# Ultra-Efficient Short Read Sequencing of T Cell Receptor Repertoires

**DOI:** 10.1101/494062

**Authors:** Janelle M. Montagne, Xuwen Alice Zheng, Iago Pinal-Fernandez, Jose C. Milisenda, Lisa Christopher-Stine, Thomas E. Lloyd, Andrew L. Mammen, H. Benjamin Larman

## Abstract

T cell receptor (TCR) repertoire sequencing is increasingly employed to characterize adaptive immune responses. However, current TCR sequencing methodologies are complex and expensive, limiting the scale of feasible studies. Here we present Framework Region 3 AmplifiKation sequencing (FR3AK-seq), a simplified multiplex PCR-based approach for the ultra-efficient analysis of TCR complementarity determining region 3 (CDR3) repertoires. By using minimal primer sets targeting a conserved region adjacent to CDR3, undistorted amplicons are analyzed via short read, single-end sequencing. We find that FR3AK-seq is sensitive and quantitative, performing comparably to two industry standards. FR3AK-seq was utilized to quickly and inexpensively characterize the T cell infiltrates of muscle biopsies obtained from 145 patients with idiopathic inflammatory myopathies and controls. A cluster of related TCRs was identified in samples from patients with sporadic inclusion body myositis, suggesting the presence of a shared antigen-driven response. The ease and minimal cost of FR3AK-seq removes critical barriers to routine, large-scale TCR CDR3 repertoire analyses.

## Introduction

T cell receptor (TCR) repertoire analysis has emerged as a powerful tool for examining adaptive immune responses. TCR repertoires, generated by the process of V(D)J recombination, encompass the T cell clones within a given individual or sample. The unique TCR of each clone defines its antigen specificity, and TCRs can be associated with distinct cellular phenotypes and tissue occupancy. Notably, TCR repertoires encompass dynamic populations of cells that represent past and current immune exposures and reactivities.^1^ Development of next generation sequencing (NGS)-based technologies over the last decade has enabled the unprecedented analysis of TCR repertoires.^2–5^

TCR recognition requires antigen processing and presentation of peptide epitopes on autologous major histocompatibility complex (MHC) molecules. Analysis of TCR repertoires both within and between individuals can be indicative of disease status, prior infections or immunizations, and individual-specific attributes of epitope selection.^6–12^ Clonally expanded or tissue-infiltrating T cell clones provide evidence of antigen-directed immune responses; integrating TCR repertoire analyses with phenotypic measurements (e.g. single cell transcriptional profiling^13^ or flow cytometric sorting of T cell subpopulations^14, 15^) can further illuminate qualitative aspects of T cell responses (**Fig 1a**).

**Figure 1:**
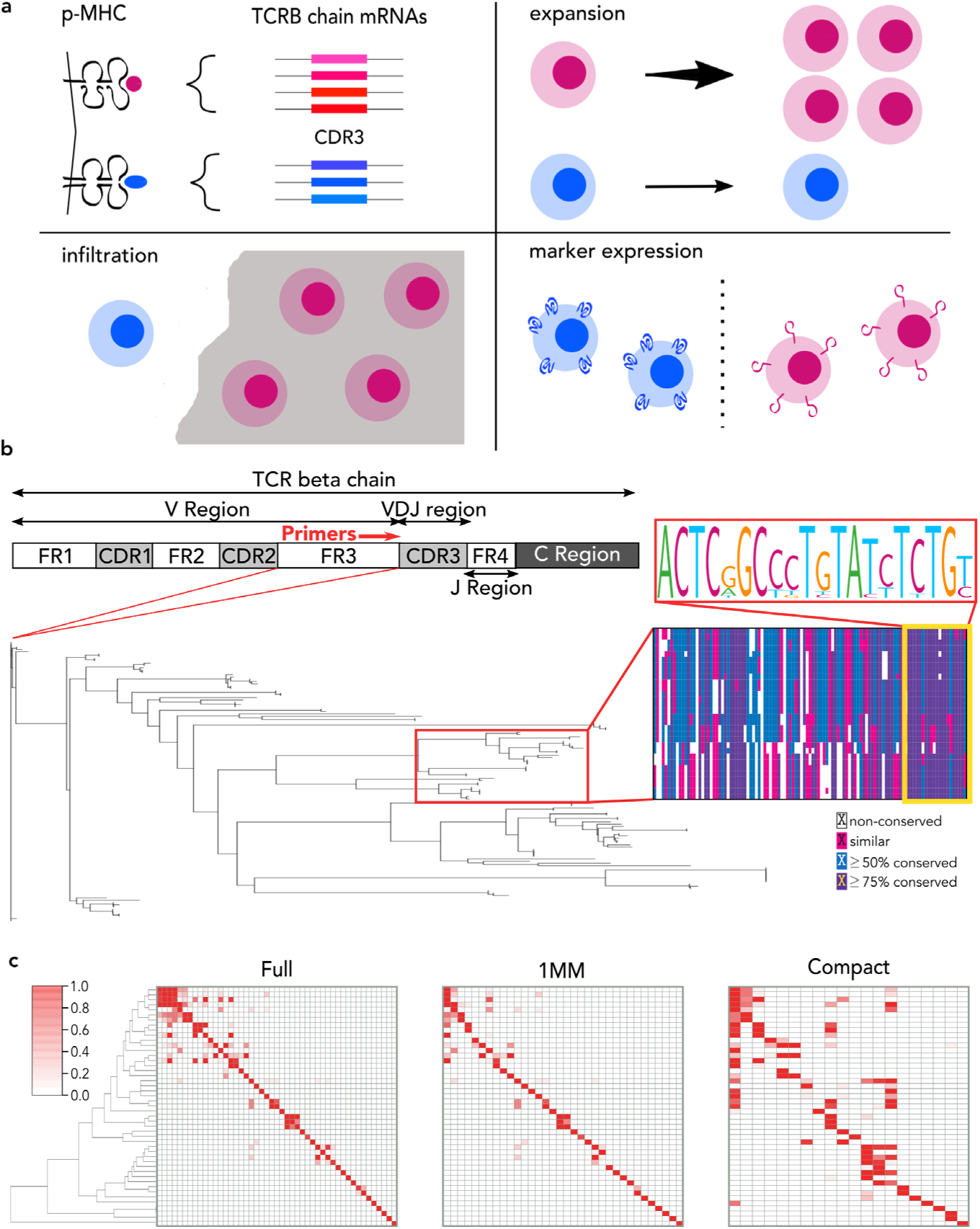
Utility of T cell receptor beta (TCRB) chain sequencing and FR3AK-seq multiplex PCR primer design. **a.** TCRB chain sequencing can be used to (i) identify related CDR3s that may share antigen specificity, (ii) detect expanded clones, (iii) identify tissue infiltrating clones, and (iv) link CDR3 sequences with T cell phenotypes. **b**. A schematic of the TCRB chain. A neighbor joining phylogenetic tree of FR3s extracted from the 128 functional human TCRBV alleles in the IMGT/GENE-DB reference directory and multiple sequence alignment from a dominant FR3 sequence cluster (red box on tree) identifies homology within the 3’ ∼20 nucleotides (yellow box). **c.** Clustered heatmaps for the Full, 1MM, and Compact primer sets showing primer amplification efficiencies for each primer across each of the 47 unique 3’ 20 nucleotides of the human TCRBV FR3 region. Inosines were considered exact matches for the Compact set. Y axis: 47 unique TCRBV FR3s. X axis: primers. The order of the primers in each heatmap can be found in **Table S1**. The order of unique TCRBV FR3s is identical to the order of the Full primers for all heatmaps.

A variety of methods have been used to generate TCR libraries for NGS, including multiplex PCR, 5’-RACE, and target enrichment.^2–5, 16, 17^ Generally, these methodologies prioritize sequencing of the hypervariable complementarity determining region 3 (CDR3) of the TCR beta (TCRB) chain. The TCRB CDR3 harbors the greatest amount of sequence diversity, confers most of the antigen specificity to a TCR, and can be used as a unique surrogate for T cell clonal identity.^18–20^ Despite their utility, however, current repertoire sequencing approaches remain complex and expensive. These limitations have greatly hindered the utilization of TCR repertoire-based analyses, particularly for studies involving large numbers of samples. We therefore sought to develop a simple, quantitative, multiplex PCR-based approach that prioritizes the ultra-efficient analysis of TCRB CDR3 sequences.

The sequence diversity of the TCRB chain variable domain, which in humans is encoded by >120 unique TCRBV alleles, necessitates the use of complex primer pools for multiplex PCR amplification. Primer competition and differential amplification efficiencies distort clonal amplicon abundance, confounding quantification.^21^ Sophisticated techniques have been developed to computationally correct amplification bias, such as the use of spike-in library standards or the incorporation of unique molecular identifiers (UMIs).^22–24^ Adaptive Biotechnologies has become a current industry leader by offering a spike-in standard corrected, multiplex PCR-based assay. At a cost of $500-$1100 per sample, however, the scale of feasible studies using this assay has been greatly constrained. Unique molecular identifier (UMI) based approaches, including the Immunoverse assay (ArcherDX), provide quantitation and enhanced sequence accuracy by resolving amplification and sequencing errors.^23–28^ However, UMI approaches typically require multi-day protocols, complicated analytical pipelines, and deeper sequencing to obtain sufficient sampling of distorted libraries.^17, 29^ By designing maximally compact primer sets and a streamlined workflow, we have essentially eliminated PCR amplification bias while maintaining high sequence level accuracy. Lower sequencing depth requirements, coupled with CDR3 analysis via single-end, short-read (100 nucleotide) sequencing, dramatically reduces the cost associated with this approach to TCR repertoire analysis.

Here, we benchmark our simplified multiplex PCR-based assay, which we call Framework Region 3 AmplifiKation Sequencing (FR3AK-seq), against two different industry standards for quantitative TCR repertoire sequencing: Adaptive Biotechnologies’ multiplex PCR-based hsTCRB immunoSEQ assay and ArcherDX’s UMI-based Immunoverse HS TCR assay. We found that FR3AK-seq data, which was analyzed using open-source software, was in excellent agreement with both. Finally, we illustrate the utility of FR3AK-seq to characterize T cell responses within biopsied muscle tissue from a cohort of 145 patients with idiopathic inflammatory myopathies and controls.

## Results

### Minimal primer sets targeting the TCRB framework region 3 (FR3) efficiently amplify CDR3 sequences

We hypothesized that minimizing the number of potentially competing V-gene primers, while also minimizing amplicon length, would maximize the efficiency of TCRB CDR3 region amplification. To identify candidate primer binding sequences upstream and proximal to CDR3, we constructed a phylogenetic tree of all functional TCRB FR3 regions available from the Immunogenetics (IMGT/GENE-DB) reference directory.^30^ This revealed a high degree of sequence conservation within the 3’ terminus of FR3 (**Fig 1b**). This homology may reflect a critical role for these sequences in the optimal presentation of the CDR3 loop. The 3’ 20 nucleotides of these IMGT FR3s were therefore extracted and used as target primer binding sequences. Remarkably, the 128 functional TCRBV alleles reported by IMGT are represented by only 47 distinct 3’-terminal 20-mers. Beyond their sequence homology, additional advantages of targeting these sequences include minimal CDR3 amplicon length (and in turn optimal amplification efficiency) and the ability to sequence using short (100 nucleotide) single end reads initiated proximal and upstream of CDR3. We refer to this approach to TCRB CDR3 analysis as FR3 AmplifiKation sequencing, or “FR3AK-seq”.

We developed a greedy algorithm to automate the design of three candidate sets of primers, each targeting the 47 distinct TCRB FR3 3’ 20-mers (**Fig S1**): one primer set lacks any universal bases or mismatches (“Full” set), one primer set lacks universal bases but allows a single mismatch to occur (“1MM” set), and one primer set contains parsimonious incorporation of up to 3 universal (inosine) bases and allows one mismatch (“Compact” set). No mismatches were permitted within the five nucleotides of the 3’ end of the primers, as a precaution to minimize interference with polymerase extension.^31–33^ The Full set consists of 47 primers (equal to the number of unique FR3 3’ 20-mers), the 1MM set consists of 34 primers, and the Compact set consists of 20 primers (**Table S1**). We calculated the expected efficiency of each primer to amplify each FR3 sequence; these efficiencies are visualized in the form of a clustered heatmap (**Fig 1c**). Importantly, each of the 47 unique 3’ FR3 20 mers are predicted to be amplified efficiently by at least one primer in each set.

### TCRB FR3 primers amplify CDR3 sequences with minimal bias

Each primer was tested separately and within its set for the ability to produce amplicon from peripheral blood mononuclear cell (PBMC) cDNA. Agarose gel electrophoresis revealed a PCR product at the expected size of ∼220 base pairs (**Fig S2a**). Importantly, incorporation of the inosine base did not prevent amplification by the KAPA Fast HotStart *Taq* DNA Polymerase. While lower PCR primer annealing temperature might increase off-target priming, it may also reduce unwanted bias in amplification from primers containing a single nucleotide mismatch.^34^ We therefore utilized gradient PCR to assess the effect of annealing temperature on the number of unique CDR3 sequences detected using the Compact primer set. As expected, more unique CDR3 clones were recovered at lower annealing temperatures (**Fig S2b**), and a similar trend was observed for the 1MM primer set. The annealing temperature that produced the maximum number of unique clones was identified for each set as follows: Full: 52°C, 1MM: 47°C, Compact: 42°C. Use of a hemi-nested RT-PCR strategy (**Fig S3**) resulted in no apparent amplification of non-TCRB sequences, even at the lowest PCR annealing temperature.

We next quantified PCR amplification bias inherent to the three primer sets. cDNA first underwent 20 cycles of PCR. An aliquot of this product was diluted 2^10^-fold and then subjected to 10 more cycles of PCR; sequencing was then performed on both amplicons and the resulting data sets compared to each other. In this experiment, significant per-cycle amplification bias would manifest as discordance. However, the concordance remains high for all three primer sets, suggesting that FR3AK-seq primers negligibly bias the CDR3 repertoire during cycles of PCR amplification (ρ = 0.997 for the Full primer set and ρ = 0.998 for the 1MM primer set, **Fig 2a**; ρ = 0.997 for the Compact primer set, **Fig S2c**).

**Figure 2:**
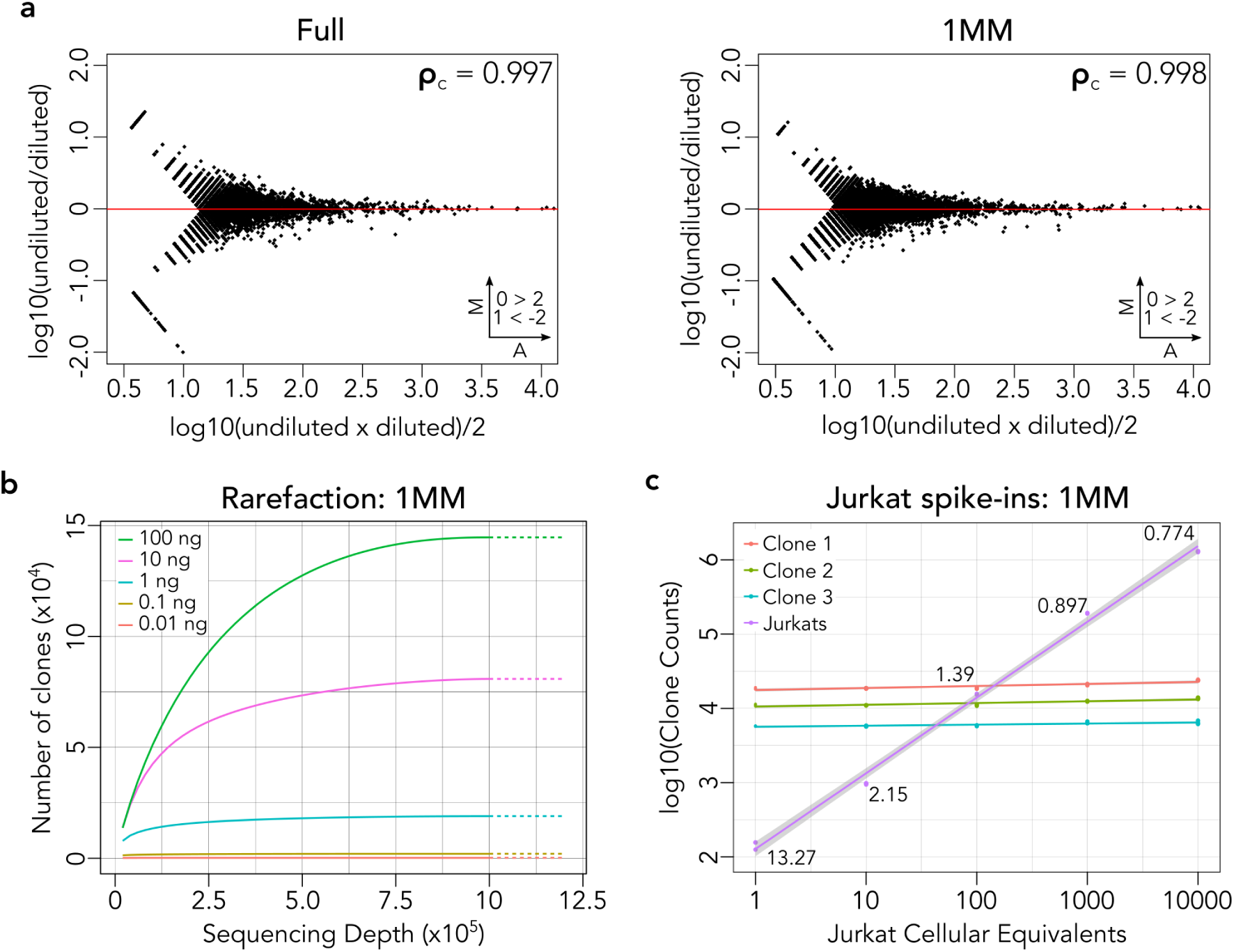
FR3AK-seq multiplex PCR performance. **a.** MA plots of 2^10^ dilution experiments for the Full and 1MM primer sets demonstrated high linear concordance (Full: 0.997, 1MM: 0.998, Compact in **Fig S2c**), indicating negligible PCR bias. Clones with ≥10 counts in either dataset were compared. Insets indicate number of clones above or below Y axis limits (set to 2 and –2 for visualization). **b.** Rarefaction analysis using T cell RNA purified from PBMCs showed the relationship between RNA input, number of clones detected, and sequencing depth using FR3AK-seq. **c.** The spiked-in Jurkat CDR3 sequence was detected at the expected abundances using FR3AK-seq (performed in triplicate). Estimated abundances of the top three T cell clones were consistent among triplicates and across spiked-in samples. 95% confidence intervals for regression lines and coefficients of variation for the cellular equivalents calculated for the T cell clones are shown.

We then performed rarefaction analysis to determine the relationship between RNA input amount, sequencing depth, and the number of unique clones detected using FR3AK-seq. We found that diversity discovery is saturated at ∼1×10^6^ reads for 100 ng of purified peripheral blood T cell RNA (**Fig 2b**), providing the rationale for RNA input amounts and sequencing depths used in subsequent analyses. We also assessed the sensitivity of FR3AK-seq to detect rare clones by spiking varying amounts of monoclonal Jurkat RNA (from 1 up to 10,000 cellular equivalents) into a background of 400 ng PBMC RNA (∼20% T cells). We found that FR3AK-seq detected the Jurkat CDR3 sequence at all input amounts at the expected abundances, while the abundance of the top three PBMC T cell clones remained constant across the dilution series (**Fig 2c**). The Jurkat sequence abundance was used to calculate cellular equivalents for the top three PBMC clones in each sample. Notably, the coefficient of variation for these cellular equivalents was below 1% in the 1,000 Jurkat cell equivalent spike-in samples (CV = 0.897%). This provided the basis for the subsequent use of 1,000 cellular equivalents of spike-in cells.

### FR3AK-seq performs comparably to a multiplex PCR-based industry standard

T cell RNA from two donors, A and B, as well as a third sample comprised of 90% Donor A RNA and 10% Donor B RNA (sample “C”), were used to generate cDNA for benchmarking studies against the current multiplex PCR-based industry standard, the immunoSEQ hsTCRB “Deep Resolution” sequencing service offered by Adaptive Biotechnologies. Separate cDNA aliquots were also subjected to FR3AK-seq analysis using each of the three primer sets. The open source software MiXCR v2.1.11 was used to define and quantify CDR3 sequences from both assays for direct comparison.^35^ The technical reproducibility of the immunoSEQ assay was quantified by comparing the abundances of Donor A’s unique CDR3 sequences against their corresponding abundances in sample C (concordance correlation coefficient, ρ = 0.989, left panel **Fig 3a**). The same analysis was applied to the FR3AK-seq data sets, which were generated using each of the three FR3 primer sets. Equally high measures of internal concordance were observed (ρ = 0.990 for the 1MM primer set, right panel **Fig 3a;** ρ = 0.990 for the Full primer set and ρ = 0.987 for the Compact primer set, **Fig S4a**). Importantly, the concordance between immunoSEQ and FR3AK-seq measurements of CDR3 abundance was also very high (ρ = 0.975, **Fig 3b**, **Fig S4b**), providing further evidence that FR3AK-seq amplifies CDR3s with negligible bias. The majority of even moderately abundant immunoSEQ-defined CDR3 sequences were also detected using any of the three FR3 primer sets (**Fig 3c**). Additionally, the 9:1 mixture composed of cells from Donor A and Donor B (sample C) was used to determine quantification accuracy of the FR3 primer sets relative to the immunoSEQ assay (**Fig 3d**, **Fig S4c**). These data indicate that all three FR3 primer sets provide CDR3 quantification comparable to the industry standard immunoSEQ assay, with the 1MM primer set performing most accurately.

**Figure 3:**
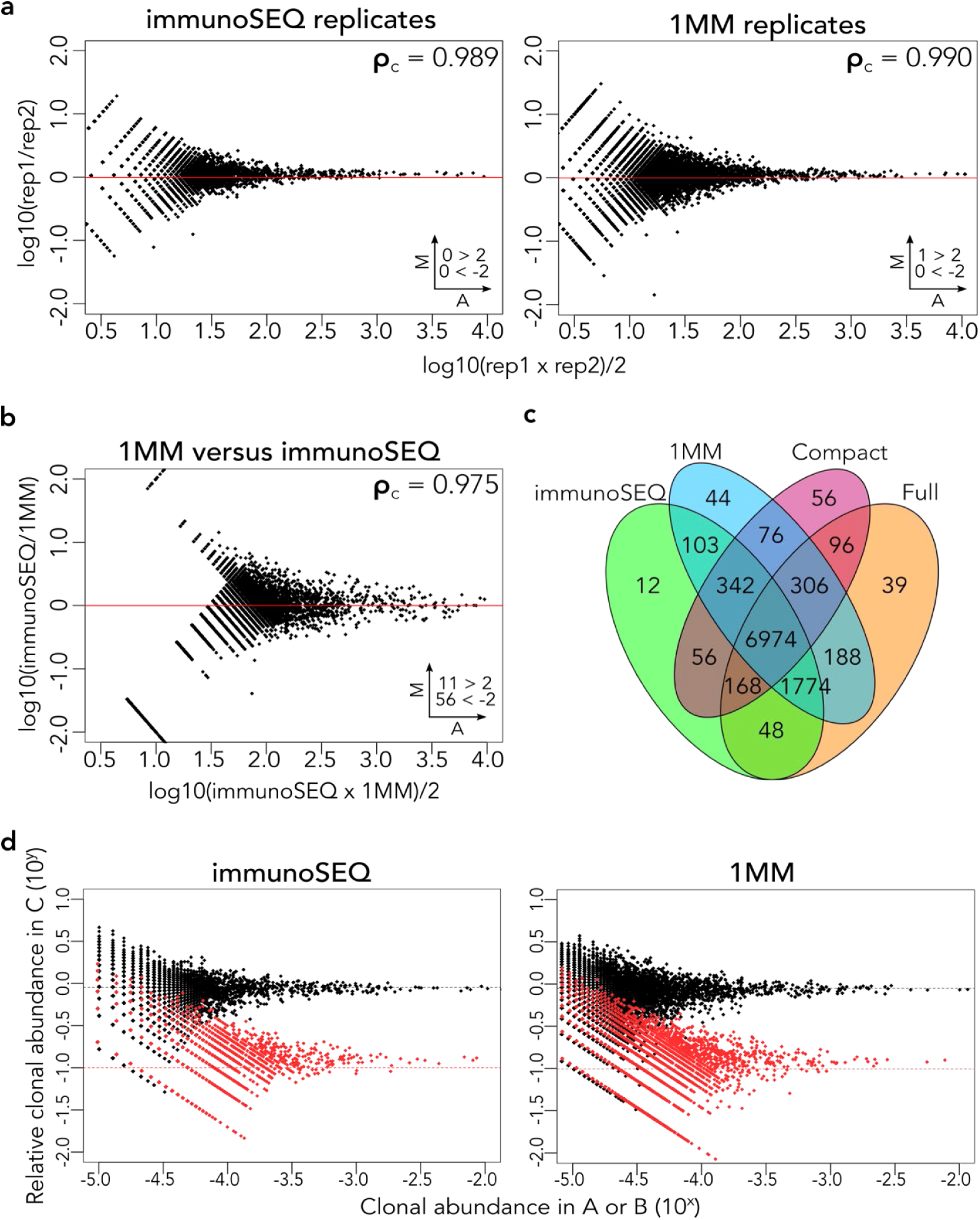
FR3AK-seq performs comparably to the multiplex PCR-based immunoSEQ assay from Adaptive Biotechnologies. **a.** MA plots comparing technical replicates of the immunoSEQ and 1MM FR3AK-seq assays. **b.** MA plot comparing the 1MM FR3AK-seq assay against the immunoSEQ assay. Clones with ≥10 counts in either dataset were included. **c.** A Venn diagram shows overlapping and non-overlapping CDR3 sequences detected using immunoSEQ and all three FR3AK-seq primer sets. CDR3 sequences were included in this analysis if they had a clone count ≥10 in any of the four data sets. **d.** Relative frequencies of Donor A (black points) and Donor B (red points) in sample C determined using the immunoSEQ assay or the 1MM FR3AK-seq assay. Insets in **a** and **b** indicate number of clones above or below Y axis limits (set to 2 and −2 for visualization).

### FR3AK-seq performs comparably to a unique molecular identifier (UMI)-based industry standard

To assess whether FR3AK-seq achieves results comparable to UMI-based methodologies, we compared the FR3AK-seq 1MM and immunoSEQ datasets to one obtained using ArcherDX’s UMI-based Immunoverse HS TCR assay. PBMC RNA from the same samples A, B, and C described above were subjected to Immunoverse analysis for comparison with the FR3AK-seq and immunoSEQ datasets. The technical reproducibility of the Immunoverse assay was similar to that of both immunoSEQ and FR3AK-seq (ρ = 0.967, **Fig 4a**). Additionally, concordance between FR3AK-seq and Immunoverse was high (ρ = 0.900, upper panel **Fig 4b**). Relatively high concordance was also observed when comparing immunoSEQ to Immunoverse (ρ = 0.886, lower panel **Fig 4b**). Crucially, the majority of clones detected by the Immunoverse assay were also detected by both immunoSEQ and FR3AK-seq (**Fig 4c**). These data demonstrate that FR3AK-seq has a high sensitivity for detecting clones of the correct sequence (4352/4403, 98.8% of total clones detected by Immunoverse). Strikingly, a large number (2803) of clones were detected by both immunoSEQ and FR3AK-seq but not Immunoverse. Rarefaction analysis of the FR3AK-seq and Immunoverse datasets explained this difference in clonal detection sensitivity, in that FR3AK-seq sampled more clones at all sequencing depths (**Fig 4d**). Although FR3AK-seq achieves better coverage of the repertoire than Immunoverse across sequencing depths, we wanted to quantify how many of the FR3AK-seq detected clones may be the result of false discovery. To address this question, we performed FR3AK-seq 1MM analysis of 1,000 cellular equivalents of Jurkat cell RNA in triplicate. Since the Jurkat cell line is monoclonal, the frequencies of subdominant clones, presumably generated by PCR or sequencing errors, can be used to assess the rate of false positive discovery. 99.98% of MiXCR-defined CDR3s mapped to the correct reported Jurkat TCRB sequence in each sample. We found an average of 50 non-Jurkat clones across triplicates, of which only one was moderately abundant (≥10 clone counts) in each. This secondary clone was the same between replicates and represented on average 0.0067% of total MiXCR-defined CDR3s. Using default MiXCR settings, FR3AK-seq analysis is therefore associated with a very low rate of false discovery due to PCR or sequencing errors. Modification of MiXCR thresholds can be used to adjust the stringency with which these subdominant clones are grouped. We also compared the CDR3 lengths of clones detected using each sequencing platform, and found that FR3AK-seq’s short (100 nucleotide) reads did not bias the distribution of CDR3 lengths recovered (**Fig 4e**).

**Figure 4:**
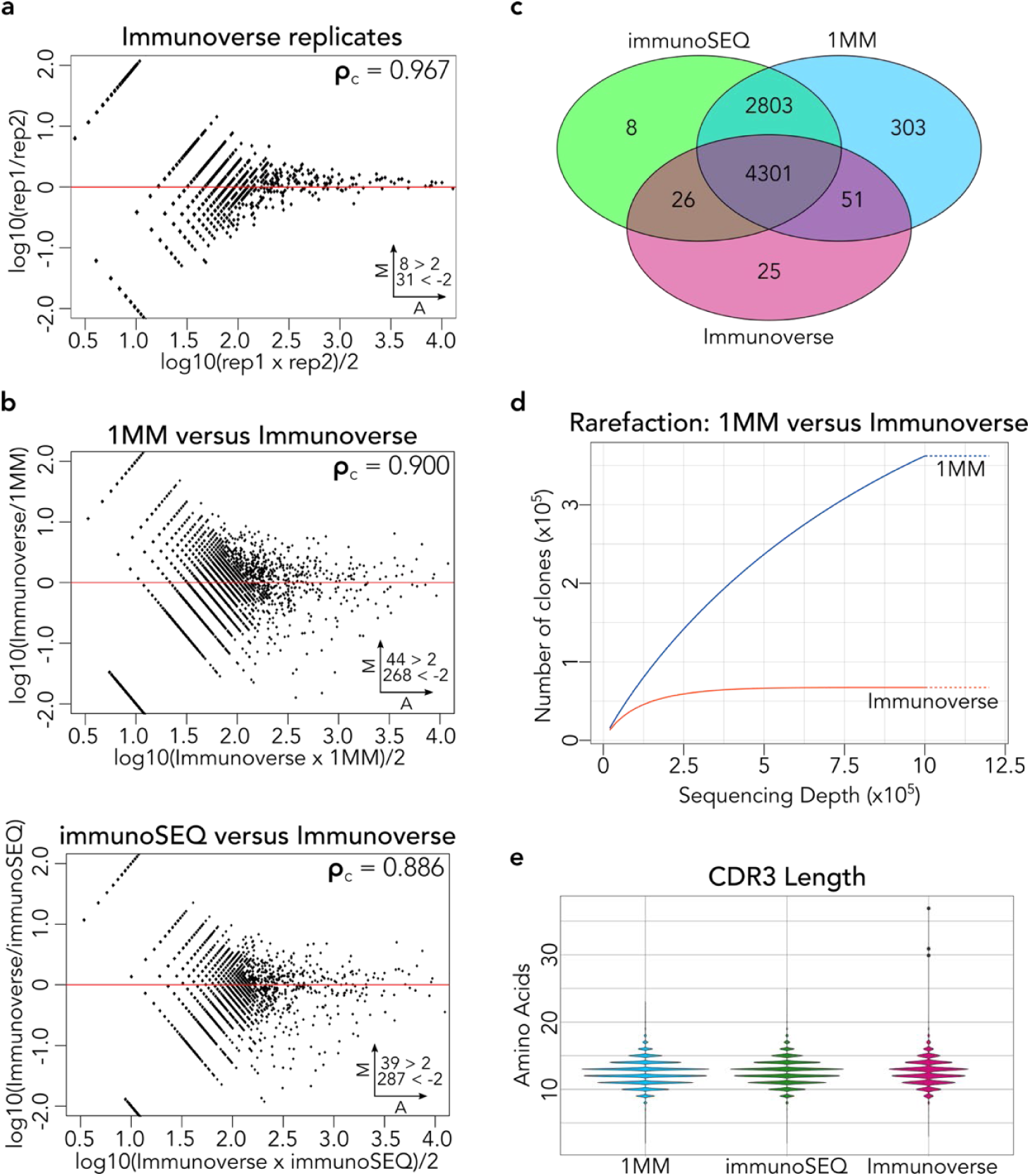
FR3AK-seq performs comparably to the UMI-based ArcherDX Immunoverse assay. **a.** MA plot comparing Immunoverse technical replicates. **b.** MA plots comparing the 1MM FR3AK-seq assay against the Immunoverse assay (top panel) and Adaptive Biotechnologies’ immunoSEQ assay against the Immunoverse assay (bottom panel). Clones with ≥10 counts in either dataset were included (using pre-deduplicated counts for the Immunoverse dataset). **c.** A Venn diagram shows overlapping and non-overlapping CDR3 sequences detected using Immunoverse, immunoSEQ, and the 1MM FR3AK-seq primer set. CDR3 sequences were included in this analysis if they had a clone count of ≥10 in any of the three data sets. **d.** Rarefaction analysis comparing the 1MM FR3AK-seq assay to the Immunoverse assay using 100 ng of T cell RNA (∼250,000 cells) for 1MM FR3AK-seq analysis and 200 ng of T cell RNA (∼500,000 cells) for Immunoverse analysis. 1MM: ∼1.2×10^6^ reads, Immunoverse: ∼1.4×10^6^ reads. **e.** Violin plots comparing CDR3 lengths captured by the short-read 1MM FR3AK-seq versus longer-read immunoSEQ and Immunoverse assays. Insets in **a** and **b** indicate number of clones above or below Y axis limits (set to 2 and −2 for visualization).

### FR3AK-seq enables inexpensive cohort-scale repertoire studies

Muscle biopsies from 145 patients with idiopathic inflammatory myopathies (IIMs, 124) as well as healthy controls (9) and non-IIM controls (12) were analyzed using the 1MM primer set at a total cost of ∼$20 per sample (**Table S2**). IIM patients included those with dermatomyositis (DM, 40), immune-mediated necrotizing myopathy (IMNM, 49), inclusion body myositis (IBM, 14), and anti-synthetase syndrome (ASyS, 21) (**Table S3**). For absolute T cell number estimation, 1,000 cell equivalents of RNA derived from a clonal tumor infiltrating T cell line^36^ were spiked into each sample after RNA purification. Aggregated CD4 and CD8 sequencing reads from RNA-seq analysis of the same samples (performed separately and reported elsewhere) tightly correlated with FR3AK-seq based estimates of cellular equivalents (**Fig 5a**). In subsequent analyses, CDR3 sequences present at levels at or below one cell equivalent per biopsy were considered ‘bystander’ T cell clones, unlikely to be involved in the disease process. By this metric, the number of non-bystander T cells were elevated in each disease subgroup versus the controls (**Fig 5b**). In agreement with previous immunohistochemical analyses, IBM biopsies contained particularly high levels of T cell infiltrates.^37–39^ We determined this to be the case both in terms of absolute T cell number and clonal diversity (**Fig 5b, Fig S5a**). No distinct patterns of J gene usage were found among patient subgroups (**Fig S6a**), and CDR3 hydrophobicity and lengths were similar between patient subgroups (**Fig S6b-c**).

**Figure 5:**
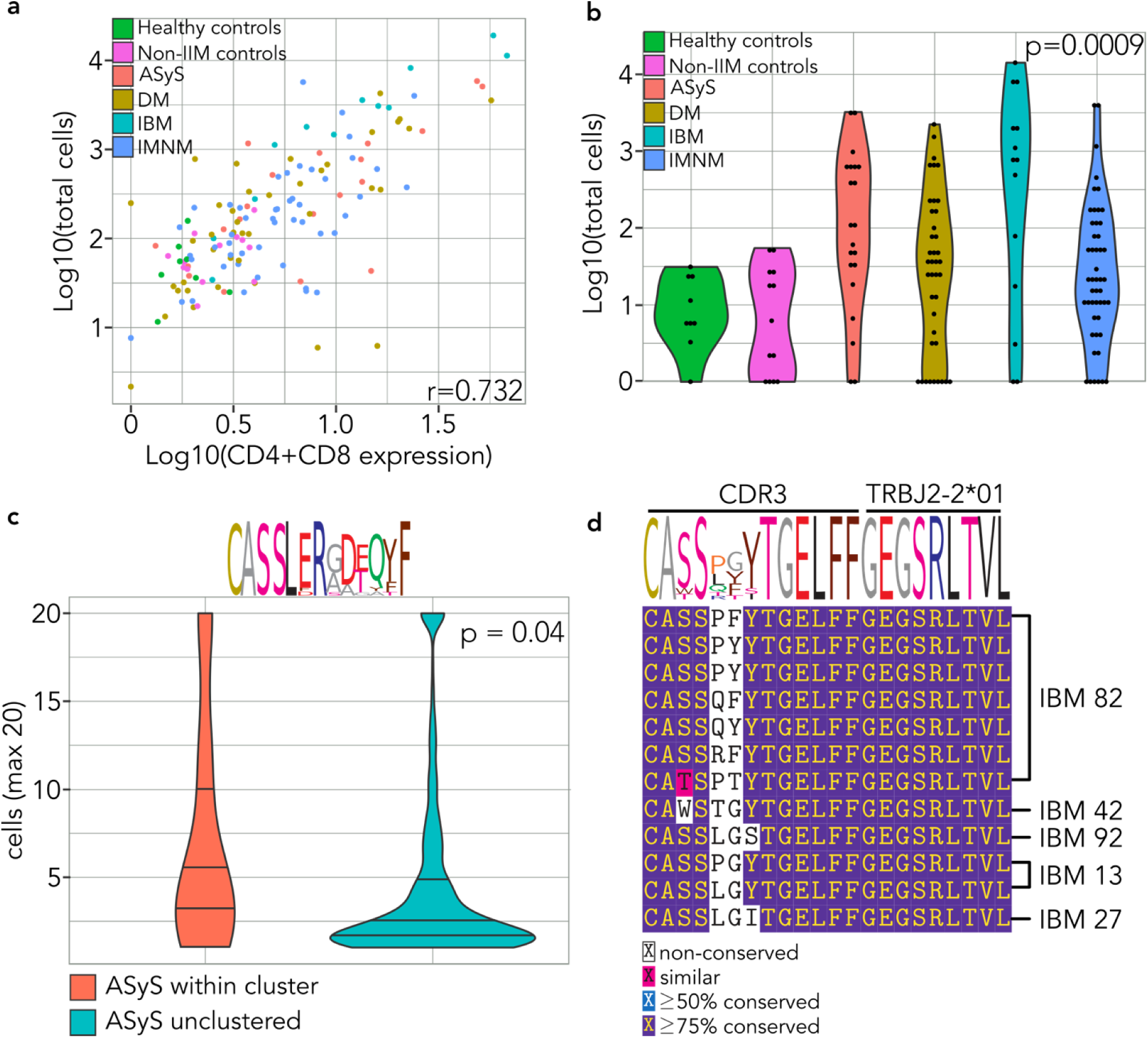
FR3AK-seq detects related T cell responses in subsets of patients with idiopathic inflammatory myopathies. **a.** Scatterplot of estimated T cell counts per patient versus CD4 and CD8 expression levels obtained from bulk mRNA-seq. Each patient data point is colored according to their IIM association. Pearson correlation coefficient (r) is shown. **b.** Distributions of cell counts per patient for each IIM and controls. Kruskal-Wallis p-value is provided. **c.** A CDR3 cluster enriched for sequences from a single patient with ASyS was identified using GLIPH software (p = 1.4×10^-7^). The amino acid sequence logo of the CDR3 region is shown above the violin plot. The clones within this cluster are more expanded than corresponding unclustered clones from the same patient. Mann-Whitney p = 0.04. Cell counts were set to a maximum of 20 for visualization. **d.** An IBM-exclusive cluster was also identified using GLIPH, encompassing 5 of 12 IBM patients who had non-bystander T cell infiltrates (41.7%, p=1.4×10^-6^). Multiple sequence alignment (using MUSCLE) is shown for the corresponding MiXCR-defined CDR3s and the J chain allele (TRBJ2-2*01).

The open source software GLIPH (grouping of lymphocyte interactions by paratope hotspots)^10^ was next used to search for clusters of related CDR3 sequences within the aggregated set of non-bystander CDR3s (**Table S4**). 143 clusters composed of sequences contributed by at least 3 individuals were identified. We first examined clusters that had a disproportionate number of sequences contributed by single individuals. **Fig 5c** provides an example of a cluster dominated by related CDR3 sequences contributed by a single ASyS patient. Compared to CDR3 sequences from the same patient which were not in any cluster, the clonal abundances of these CDR3s were significantly increased, suggestive of a polyclonal, shared antigen-driven expansion. These results suggest that GLIPH is able to cluster related CDR3 sequences that have been detected and quantified using FR3AK-seq.

We next determined whether any of the GLIPH clusters were composed of sequences contributed disproportionately by individuals from one of the IIM subgroups. Indeed, we found a GLIPH cluster that was composed exclusively of sequences contributed by IBM patients. Specifically, 5 of the 12 IBM patients with non-bystander T cells contributed at least one sequence to this cluster (p = 1.4×10^-6^, **Fig 5d**). Remarkably, all CDR3 sequences in this IBM cluster are linked to a single J-chain (TRBJ2-2*01), providing additional support for shared antigen recognition. Taken together, this study demonstrates the utility of FR3AK-seq in probing T cell responses at cohort scale.

## Discussion

Although the utility of TCR repertoire sequencing to characterize T cell responses has been well established, the lack of a streamlined, quantitative, inexpensive, and non-proprietary assay has limited the scope and scale of feasible studies. By rationally designing minimal sets of primers that target the 3’ terminus of FR3, we have developed a multiplex PCR-based approach for ultra-efficient library preparation and sequencing of TCRB CDR3 repertoires. By minimizing amplification bias (via reduction of primer number and amplicon length), the resulting sequencing libraries quantitatively capture clonal abundance distributions with an accuracy comparable to both multiplex PCR-based and UMI-based industry standards. While the library preparation and sequencing strategy presented here already bring the per-sample cost to ∼$20 (**Table S2**), we expect that alternative approaches to library preparation, reduced sequencing depth, and declining sequencing costs will likely enable another ∼10-fold reduction in per-sample cost. In addition to reducing the overall cost of obtaining CDR3 sequences, we have developed the FR3AK-seq workflow to require minimal expertise, effort, and time. Quantitative sequencing libraries can easily be prepared within a single day. Comparisons of protocols and cost between FR3AK-seq, immunoSEQ, and Immunoverse are summarized in **Table S5**.

Our approach to TCRB repertoire analysis should be generalizable to additional immune receptor types, as well as to those of non-human species (see **Table S6** for *Mus musculus* TCRB primers). Although these mouse primers have not been as extensively validated as the human set, they have been confirmed to efficiently amplify CDR3 sequences from mouse tissues. We have additionally used the same FR3 primer design and analysis principles to target human TCR alpha, gamma, and delta repertoires, although these primers have not yet been validated (**Table S7**). We have also designed a set of TCRB J region reverse primers (**Table S8**) that perform comparably to our constant region reverse primer using cDNA templates (ρ = 0.949, **Fig S7**). These will be potentially useful for analysis of FFPE samples which suffer from extensive RNA degradation. Interestingly, during the preparation of this manuscript, the Euroclonality Group (creators of the BIOMED2 primer set^40^) published a new set of TCR primers for next-generation sequencing, including some that are designed in the 5’ portion of FR3,^41^ providing additional validation for an FR3-targeted approach to TCR repertoire sequencing.

V gene usage is often of interest, for example to detect usage skewing and for functional studies involving TCR cloning. FR3AK-seq prioritizes small amplicons to reduce both PCR bias and cost. Therefore, by design, amplicons do not contain easily recoverable V gene sequence information. However, it is theoretically possible to make inferences on each CDR3’s associated V genes, based on the pattern of FR3 primer usage. We expect these inferences to be more accurate for more abundant clones. When analysis of V gene usage is desired, complementary techniques (e.g. 5’RACE) can be parsimoniously utilized to capture this information. One may wish to use FR3AK-seq to track unique CDR3 sequences of interest over time, across tissues, and/or after FACS analysis – these FR3AK-seq detected CDR3 sequences can then be readily associated with a V gene by merging data sets.

To demonstrate the use of FR3AK-seq for efficient cohort scale analysis of TCRB CDR3 repertoires, we characterized the muscle-infiltrating T cells present within muscle biopsies obtained from 145 inflammatory muscle disease patients and controls. TCRB CDR3 sequence clustering indicated both donor and disease specific antigen-driven T cell responses. Importantly, the majority of T cell clones detected in IBM muscle biopsies were not hyperexpanded, in contrast to the hypothesis that infiltrating IBM T cells originate from a clonal T cell large granular lymphocytic leukemia (TLGLL) (**Fig S5b**).^42, 43^ Future studies will utilize FR3AK-seq in combination with *in vitro* stimulation to characterize the antigenic determinants of these clonal clusters.

## Methods

### FR3 primer design

FR3 sequences from all functional human TCRB V alleles were downloaded from the IMGT/LIGM-DB reference directory (non-functional and pseudogenes were excluded).^30^ Multiple sequence alignment (MSA) was performed using MUSCLE and UPGMA (Unweighted Pair Group Method with Arithmetic Mean) and neighbor-joining phylogenetic trees were generated via the R package msa.^44^ The 3’ 20 nucleotides of the TCRB chain FR3s were subsequently used to design three sets of primers as described in the text. We automated design of the 1MM and the Compact primer sets with an algorithm outlined in **Fig S1**, available on Github (https://github.com/jmmontagne/FR3AK-seq). Human TCRB primer sequences can be found in **Table S1**.

### PBMC preparation, T cell purification, RNA extraction, and cDNA synthesis

PBMCs from Donors A and B were isolated by Ficoll-Paque (GE Healthcare, Pittsburgh, PA) gradient centrifugation and cryopreserved. T cells were purified from thawed PBMCs using the EasySep Human T cell enrichment kit (STEMCELL Technologies, Cambridge, MA). RNA was extracted from purified human T cells using the RNeasy Plus Minikit (Qiagen, Germantown, MD). 0.5 μg Donor B RNA was mixed with 4.5 μg Donor A RNA to create sample C for quantitation experiments. 4 μg each of Donor A, Donor B, and sample C RNA was reverse transcribed using a TCRB chain constant region reverse primer with Superscript III First-Strand Synthesis System (Invitrogen, Waltham, MA). cDNA was column purified with the Oligo Clean and Concentrator Kit (Zymo Research, Irvine, CA). 100 ng of purified cDNA from each sample was sent to Adaptive Biotechnologies (Seattle, WA) for the immunoSEQ hsTCRb Deep sequencing service or used for FR3AK-seq PCR. For the ArcherDX (Boulder, CO) Immunoverse analysis, RNA was extracted from matched Donor A and B PBMCs and mixed at the same 9:1 ratio to create sample C. 800 ng of RNA from each sample was input into the Immunoverse assay. The concordance between FR3AK-seq and Immunoverse remained high despite the use of PBMC RNA rather than T cell RNA for the Immunoverse assay. All primer sequences are provided in **Table S1**.

### Polymerase Chain Reaction (PCR) and sequencing library preparation

100 ng TCRB chain cDNA was used as template for PCR with KAPA2G Fast Multiplex Mix (Kapa Biosystems, Wilmington, MA). Forward TCRB FR3 primers from the Full, 1MM, or Compact primer sets were used with a single nested TCR beta chain constant region reverse primer at 0.2 μM per primer. Samples underwent 30 cycles of PCR for visualization by agarose gel electrophoresis, or 20 cycles for NGS library preparation (“PCR1”). We compared the performance of Herculase II Fusion DNA polymerase (Agilent, Santa Clara, CA) and KAPA2G Fast Multiplex Mix for PCR1 and found that both enzymes perform comparably (ρ = 0.966, **Fig S8**). Herculase II is therefore an acceptable enzyme alternative for PCR1 if desired.

20 cycles of PCR2 were performed on PCR1 product (2 μL of PCR1 added to 18 μL of PCR2 master mix, which contained 0.25 μM each i5 and i7 sample barcoding primers) to incorporate sample barcodes and Illumina sequencing adaptors using Herculase II Fusion DNA Polymerase. Equal volumes of barcoded PCR2 products (5 μL each) were pooled and PCR column purified using QIAquick PCR Purification Kit (Qiagen). Libraries were quantified using KAPA Library Quantification Kit for Illumina Platforms (Kapa Biosystems). Barcoding primer sequences can be found in **Supplementary File 1**. See **Fig S3** for a schematic of our sequencing strategy.

### Sequencing and CDR3 analysis

Sequencing was performed on an Illumina NextSeq 500 (immunoSEQ benchmarking comparisons), MiSeq (Immunoverse samples), or HiSeq 2500 (IBM muscle biopsy analysis). CDR3s were identified and quantified using MiXCR v2.1.11 software^35^ with default parameters except as noted in **Table S9**. Data obtained from Adaptive Biotechnologies’ immunoSEQ assay were re-analyzed using MiXCR v2.1.11 and identical parameters for comparison to our own data. For reanalysis, full nucleotide sequences for each CDR3 from the Adaptive Biotechnologies ImmunoSEQ dataset were expanded to repeat as many times as indicated by the corresponding “count” column. This file, with each clonal nucleotide sequence represented as many times as its “count” column, was converted to FASTA format for compatibility with MiXCR. The sequences obtained from ArcherDX’s Immunoverse assay were expanded to repeat as many times as their pre-deduplicated clone counts and then matched back to their deduplicated clone counts after reanalysis with MiXCR v2.1.11. Clone count cutoffs for each analysis are as described in the figure legends and refer to the pre-deduplicated counts for the Immunoverse assay (Note: deduplicated clone counts were used for all comparisons, while pre-deduplicated counts were only used to filter the dataset to ≥ 10 counts). Information regarding the total number of clones, the number of singletons, and the number of clones occurring at ≥10 counts for FR3AK-seq, immunoSEQ, and Immunoverse can be found in **Table S10**.

### Repertoire analysis using muscle biopsies from patients with idiopathic inflammatory myopathies (IIMs)

RNA isolated from muscle biopsies taken from 145 IIM patients or controls was reverse transcribed, PCR amplified, sequenced, and analyzed as described above. Healthy control muscle biopsies were obtained from healthy volunteers at the Skeletal Muscle Biobank of the University of Kentucky, while non-IIM controls were obtained for clinical purposes at the Johns Hopkins Neuromuscular Pathology Laboratory and were normal after pathologic evaluation. We added 1,000 cell equivalents of RNA from a clonal tumor infiltrating T cell line as a spike-in to each sample.^36^ Sequencing of 6 spike-in only samples provided the CDR3 sequences to remove from downstream analyses of samples containing muscle biopsy RNA. The spike-in counts were used for cell number quantification within patient samples, which distinguished ‘bystander’ versus ‘non-bystander’ T cell clones. One non-IIM control muscle biopsy was sequenced in duplicate, making the total number of samples 146.

CDR3 sequences obtained from these samples were analyzed for disease-specific clusters using the GLIPH 1.0 group discovery algorithm (see **Table S9** for parameters).^10^ To determine the statistical significance of each cluster, we performed chi-square analysis on all clusters with at least three contributing individuals, followed by Benjamini-Hochberg multiple comparison correction. This analysis was performed for both the number of sequences contributed by each disease subgroup to each cluster, as well as the number of patients in each disease group represented within each cluster. P-values ≤0.05 after multiple test correction were considered statistically significant.

## Author Contributions

Conceptualization and experimental design: J.M.M., H.B.L.; performing experiments: J.M.M.; data analysis and software: J.M.M., X.A.Z.; human specimen procurement and analysis: I.P-F., J.C.M., L.C-S., T.E.L., A.L.M., H.B.L.; data interpretations: J.M.M., X.A.Z., T.E.L., A.L.M., H.B.L.; paper writing: J.M.M., H.B.L. All authors reviewed the paper.

## Competing Interests

H. Benjamin Larman is a consultant for Tscan Therapeutics, which seeks to develop T cell receptor based cellular therapies.

## Acknowledgements

The authors would like to thank Dr. Erik Wright for his advice on primer design, Dr. Suzanne L. Topalian for the TIL1558 cell line, and Dr. Maximilian F. Konig for his review of the manuscript. We would also like to thank the Johns Hopkins Medical Institute Deep Sequencing and Microarray Core Facility and the Johns Hopkins Institute of Genetic Medicine Genetics Resources Core Facility. This work was supported by NIH grant U24AI118633 and a Prostate Cancer Foundation Young Investigator Award.

## SUPPLEMENTARY FIGURES

**Figure S1:**
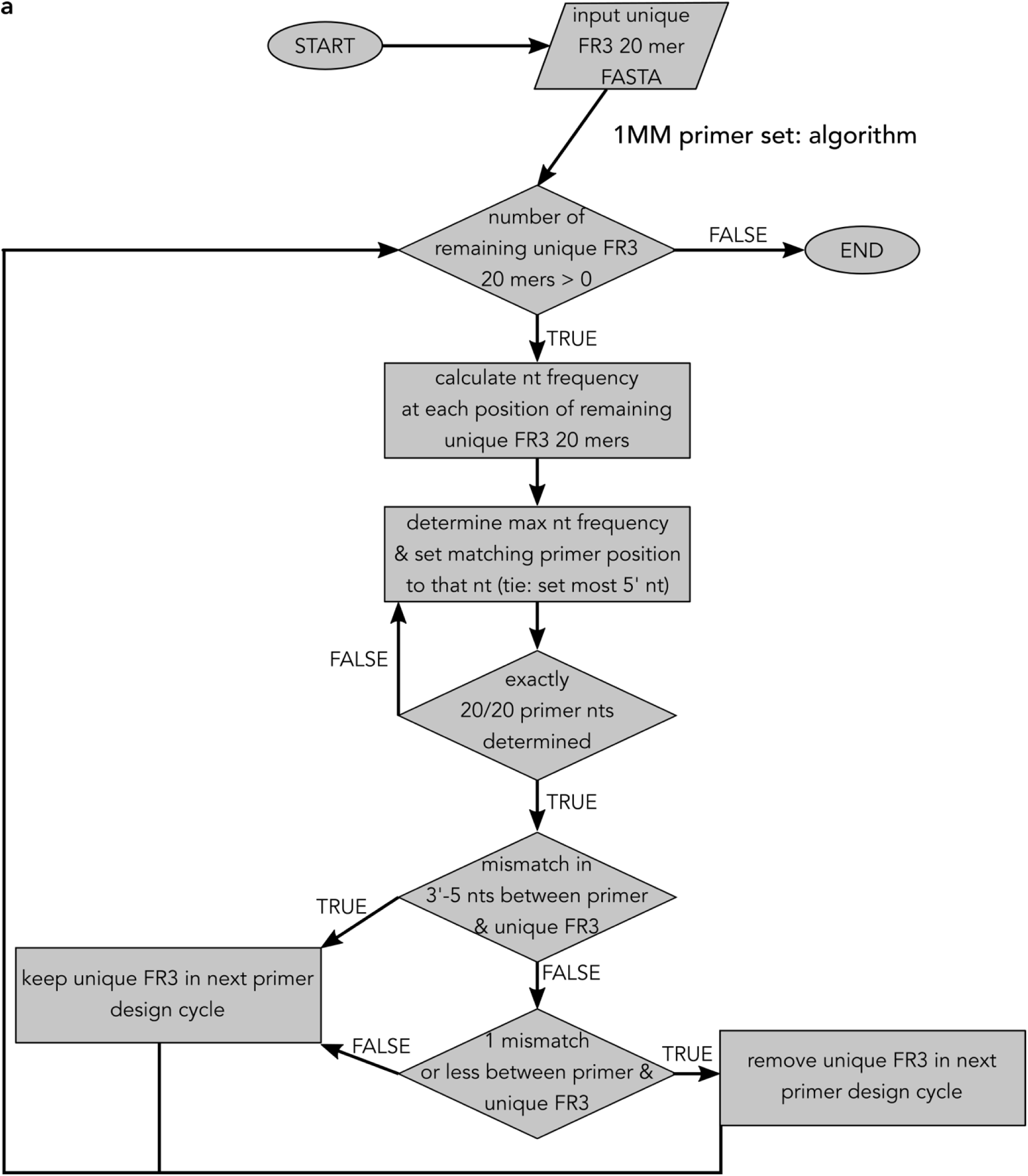
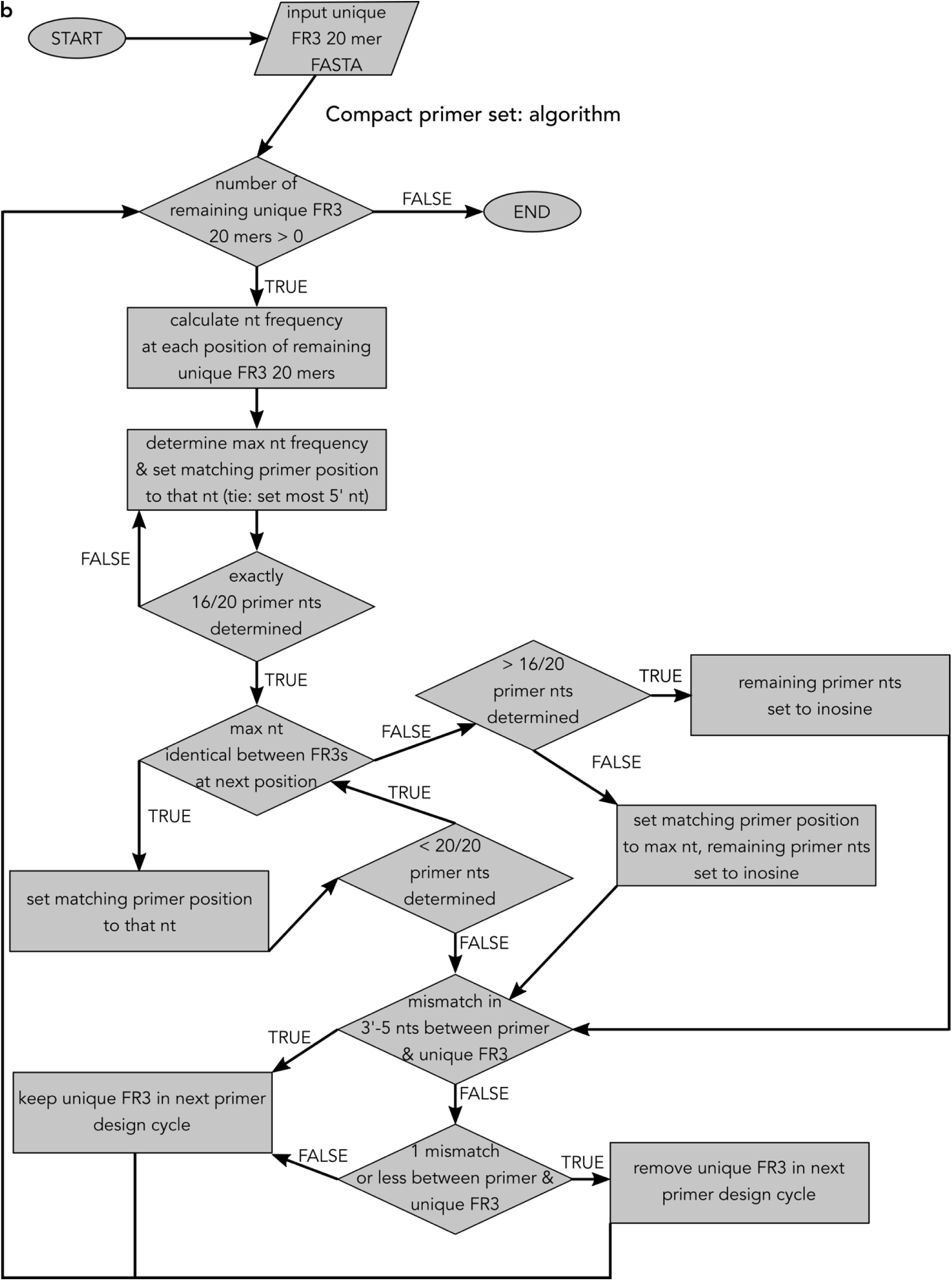
FR3AK-seq primer design algorithm schematic. **a.** 1MM primer set design algorithm. Input is a FASTA formatted file containing each unique FR3 20-mer nucleotide (nt) sequence. Nt frequencies at each of the 20 FR3 positions were calculated. The position with the maximum nt frequency in the unique FR3 20-mers was used and the corresponding position in the primer was set to that nt. In the case of ties, the most 5’ position was set first. Once all 20 nts of the primer were determined, unique FR3 20-mers were removed from the next round of primer design only if they: i) did not have a mismatch in the most 3’-5 nts (as described in the text) and ii) had 1 mismatch or an exact match to the designed primer. The primer design loop continued until no unique FR3 20-mer sequences remained. **b.** Compact primer set design algorithm. Primers were designed as in **a** until 16/20 primer nts were determined. At this point, if the position with the maximum nt frequency in the unique FR3 20-mers was identical to the number of remaining FR3 20-mers, then these nts were set to these positions. If they were not and exactly 16/20 nts in the primer were already determined, then the nt with the maximum frequency was set to the corresponding primer position and the remaining 3 nt positions in the primer were set to inosines. If they were not and >16/20 nts in the primer were already determined, then the remaining nt positions were set to inosines. In this way, we allowed for up to 3 inosines and 1 mismatch. Once all 20 nts of the primer were determined, the remaining design was identical to **a**, where we considered each inosine a match to any nt in the FR3 regardless of its position in the primer.

**Figure S2:**
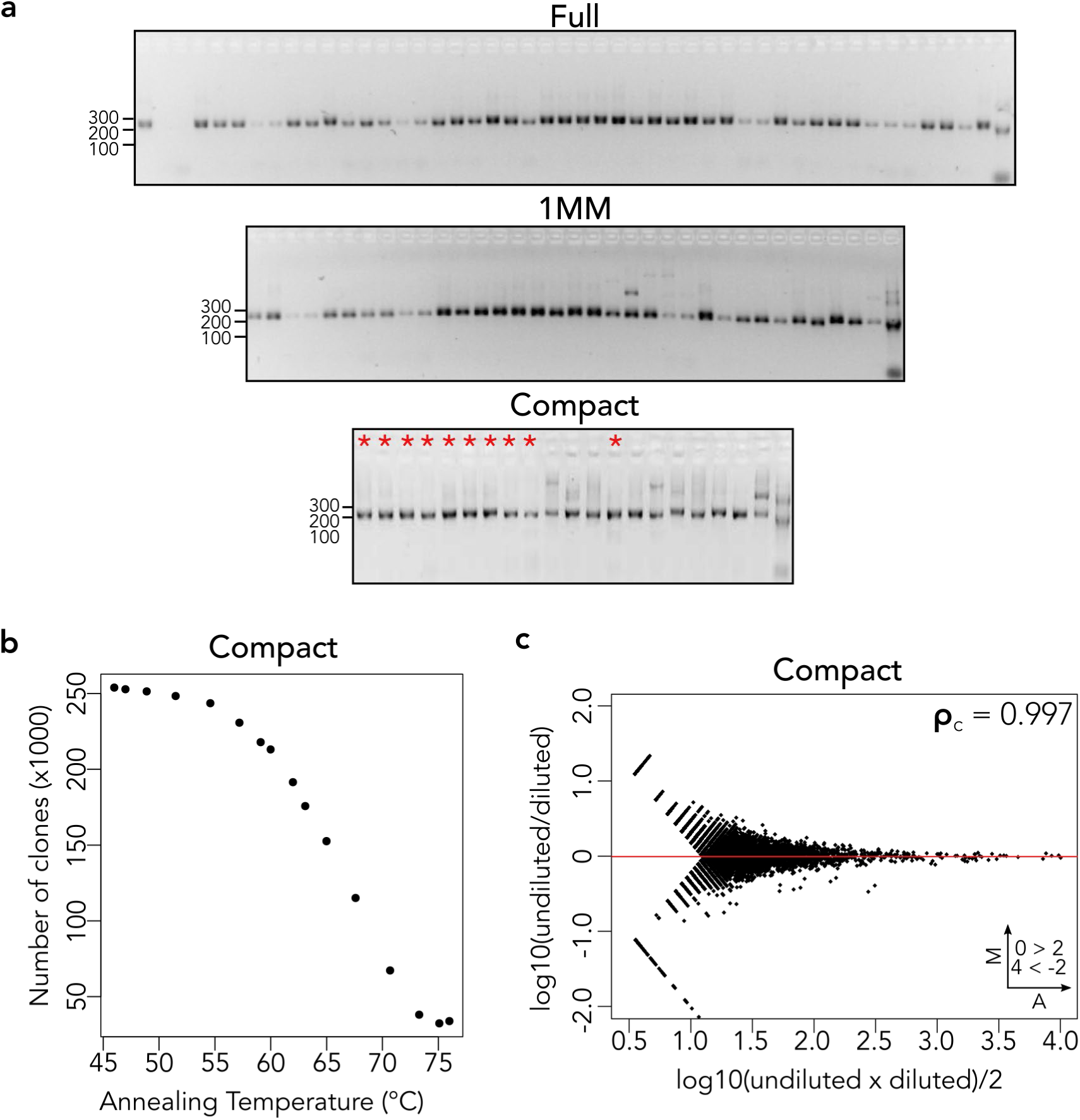
FR3AK-seq multiplex PCR performance. **a.** Individual primers from all three sets produce amplicons of the expected size from human PBMC cDNA (∼220 bp). Asterisks indicate inosine-containing primers. Order of primers can be found in **Table S1**. **b.** The number of clones detected versus annealing temperature using the Compact primer set is shown. A similar trend was observed for the 1MM primer set. To account for differences in number of reads between samples, FASTQ sequencing files were trimmed to identical line numbers before analysis. **c.** MA plot of 2^10^ dilution experiment for the Compact primer set demonstrated high linear concordance (ρ=0.997), indicating negligible PCR bias. Clones with ≥10 counts in either dataset were compared. Insets indicate number of clones above or below Y axis limits (set to 2 and –2 for visualization).

**Figure S3:**
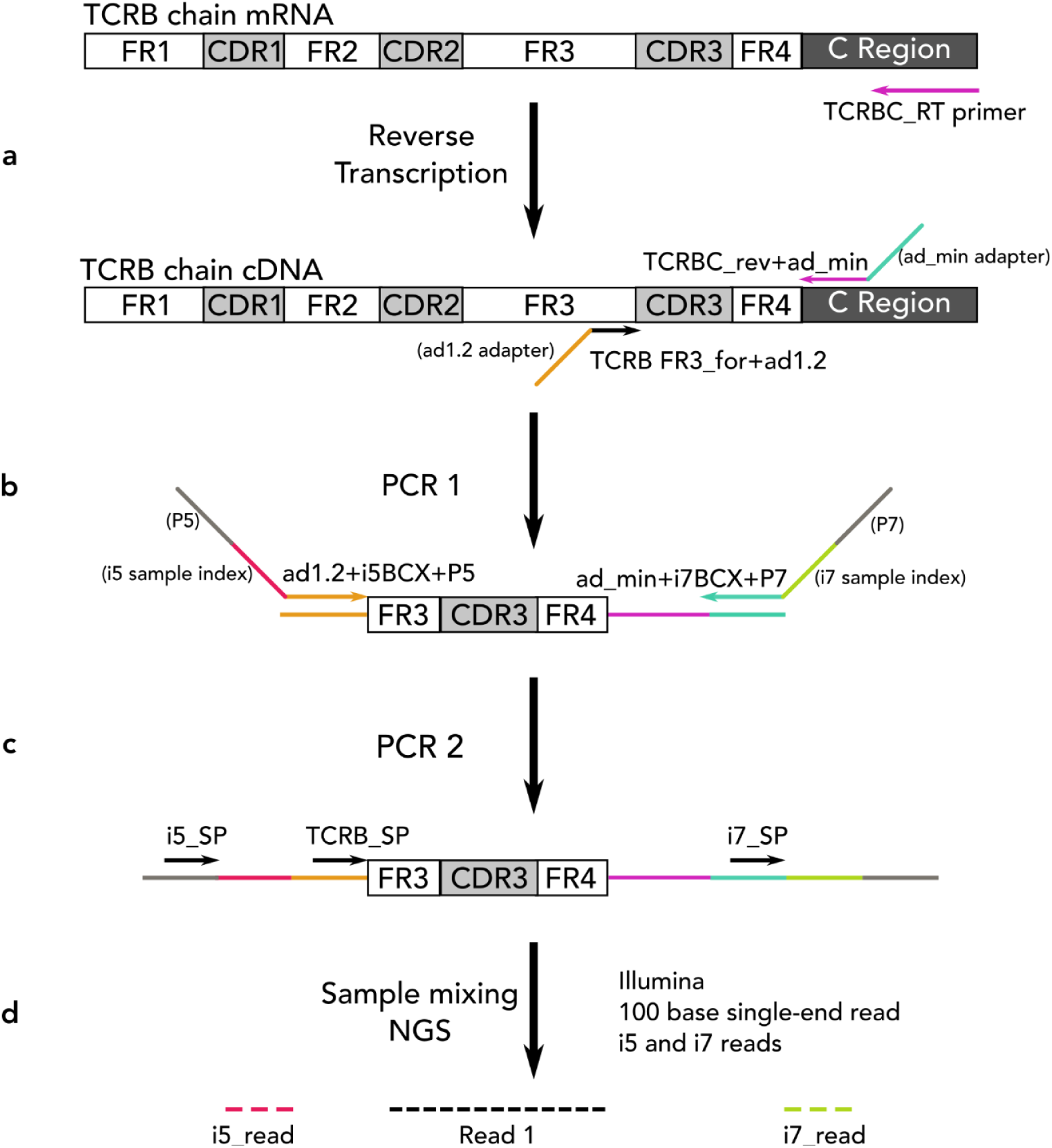
FR3AK-seq dual index sequencing strategy. **a.** Reverse transcription using a TCRB constant region primer generates cDNA for use as template in **b.** PCR1 using our forward TCRB FR3 primers and a hemi-nested TCRB constant region reverse primer. FR3 primers have 5’ ad1.2 adapter sequences and the constant region primer has a 5’ ad_min adapter sequence. **c.** PCR2 using PCR1 product as a template and indexed ad1.2 and ad_min primers, including P5 and P7 adapters for Illumina sequencing. Samples are mixed upon completion of PCR2 prior to **d.** Next Generation Sequencing (NGS): Illumina 100-base single-end reads along with index 5 (i5) and index 7 (i7) reads. Dual indexes are used for demultiplexing reads from each sample and can be found in **Supplementary File S1**. No index read mismatches were permitted during sample demultiplexing. Primer and adapter sequences can be found in **Table S1**.

**Figure S4:**
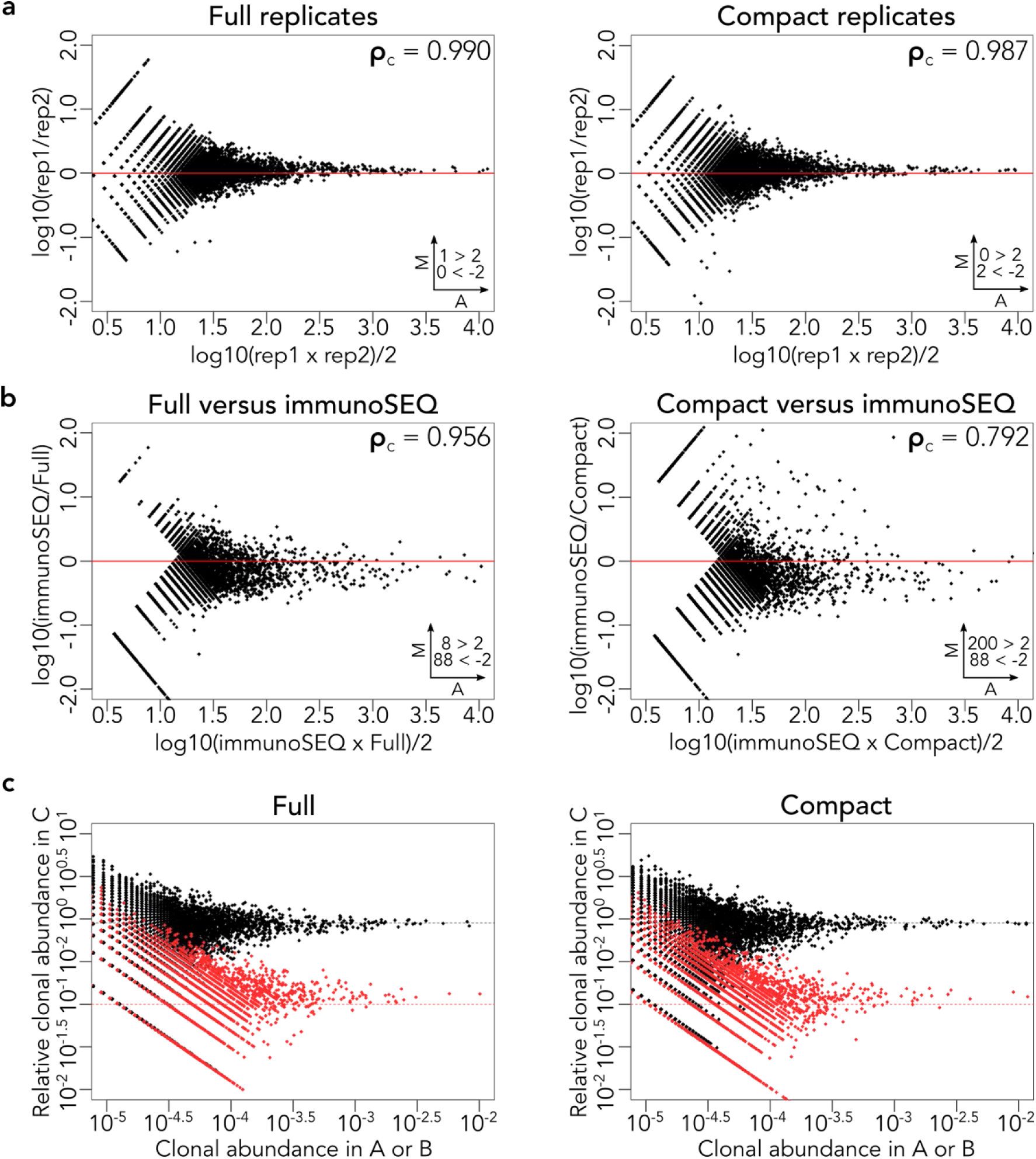
FR3AK-seq Full and Compact primer sets perform comparably to immunoSEQ. **a.** MA plots showing technical replicates of the Full (left panel) and Compact (right panel) FR3AK-seq assays. **b.** MA plot comparing the Full (left panel) and Compact (right panel) FR3AK-seq assay against the immunoSEQ assay. Clones with ≥10 counts in either dataset were included. **c.** Relative frequencies of Donor A (black points) and Donor B (red points) in sample C determined using the Full (left panel) and Compact (right panel) FR3AK-seq assay. Insets in **a** and **b** indicate number of clones above or below Y axis limits (set to 2 and –2 for visualization)

**Figure S5:**
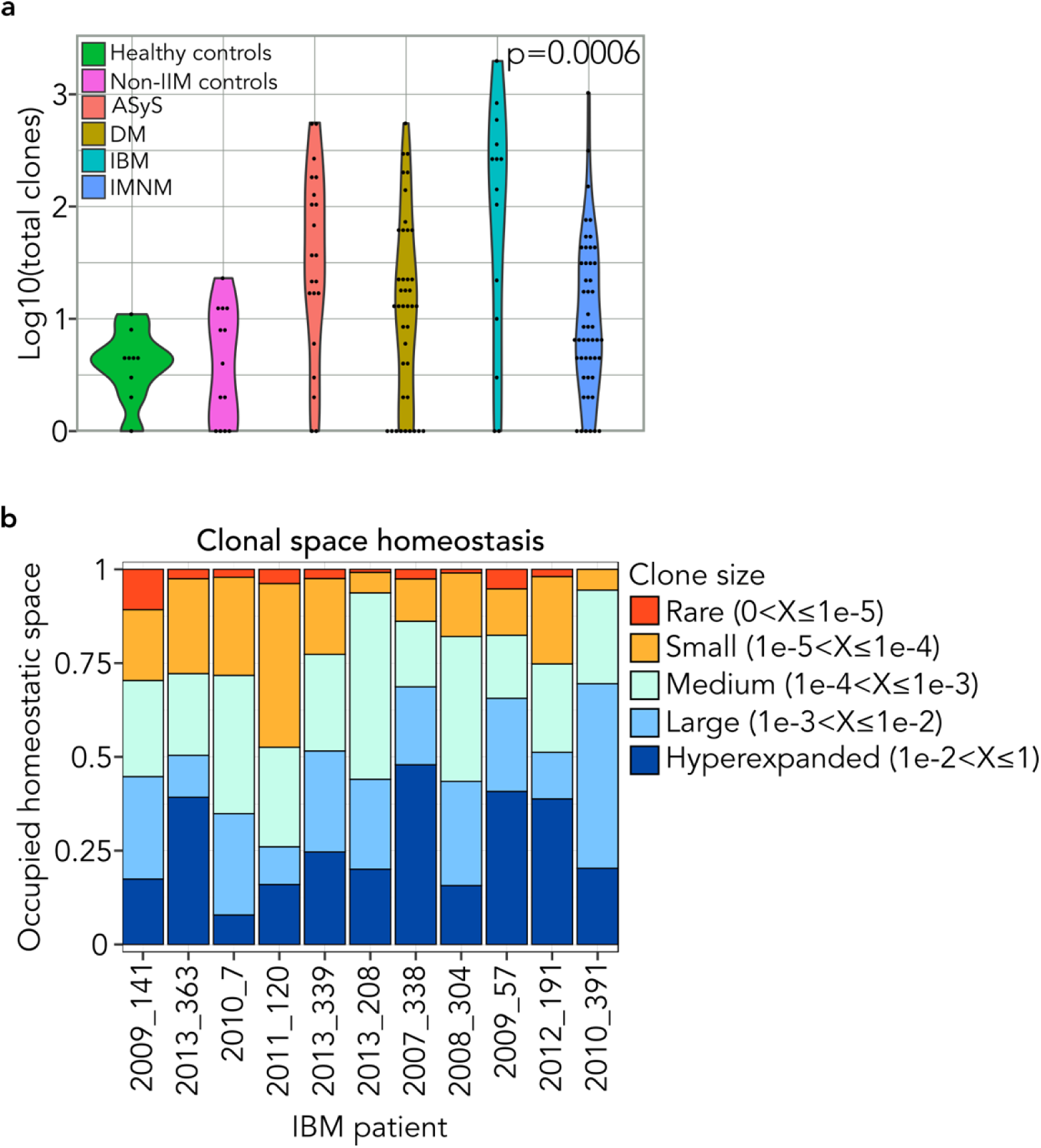
a. Distributions of unique clones per patient for each IIM subgroup and controls. Kruskal-Wallis p-value is shown. **b.** Clonal space homeostasis for each IBM donor with at least one clone present at >2 T cell equivalents in their muscle biopsy. Criteria for clone size are shown.

**Figure S6:**
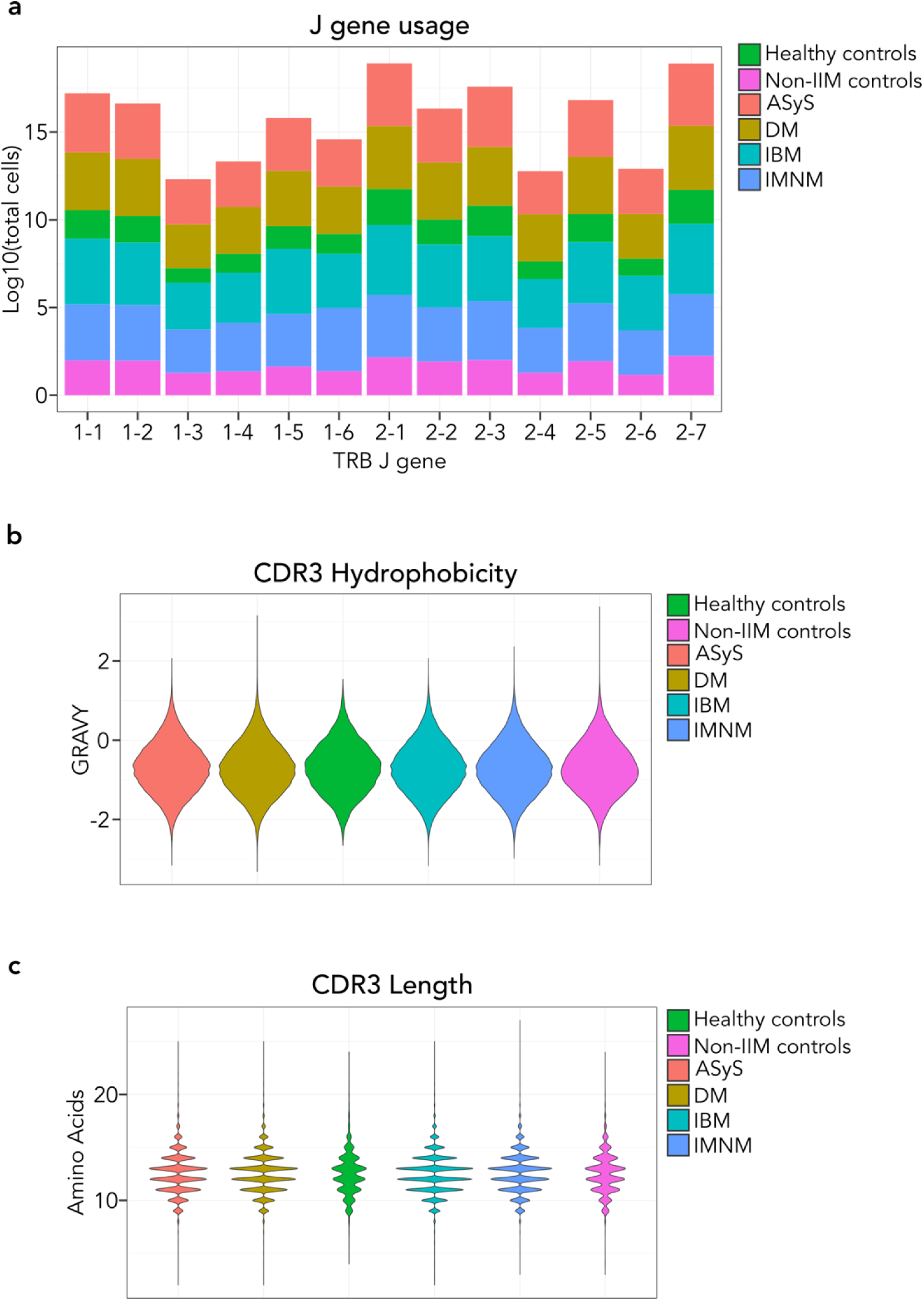
Summary statistics of FR3AK-seq TCRB repertoires from each IIM subgroup and controls. **a.** Stacked bar plot of J gene usage for each IIM subgroup and controls. **b.** Grand Average of Hydropathy (GRAVY) for CDR3s from each IIM subgroup and controls. **c.** CDR3 length comparisons between each IIM subgroup and controls.

**Figure S7:**
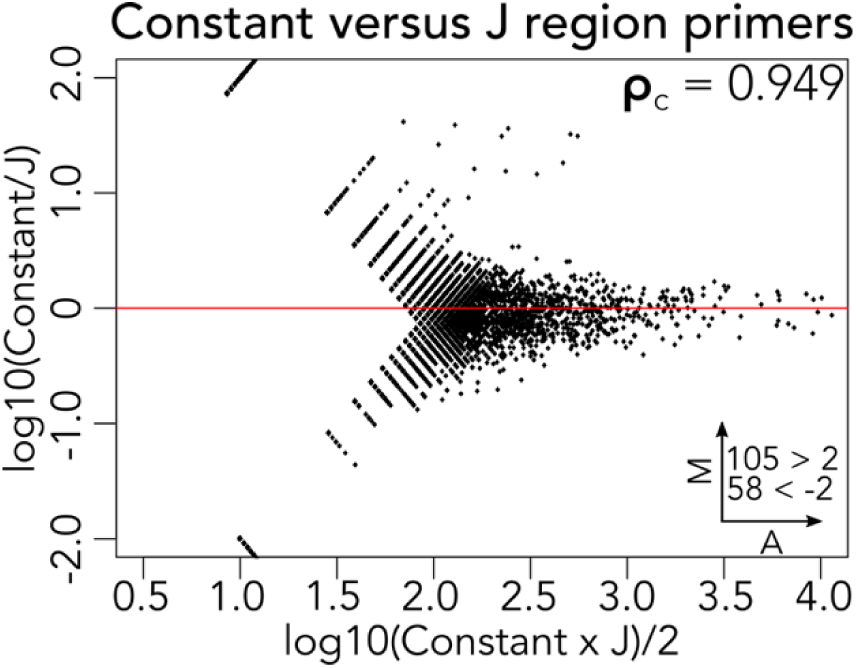
MA plot demonstrating the performance of the FR3AK-seq constant region reverse PCR1 primer as compared to a set of J region reverse primers. cDNA was used as a starting material along with the 1MM forward primer set with PCR conditions as described in the Methods section. CDR3 sequences were included in this analysis if they had a clone count of ≥10 in either data set. J region primers can be found in **Table S8**.

**Figure S8:**
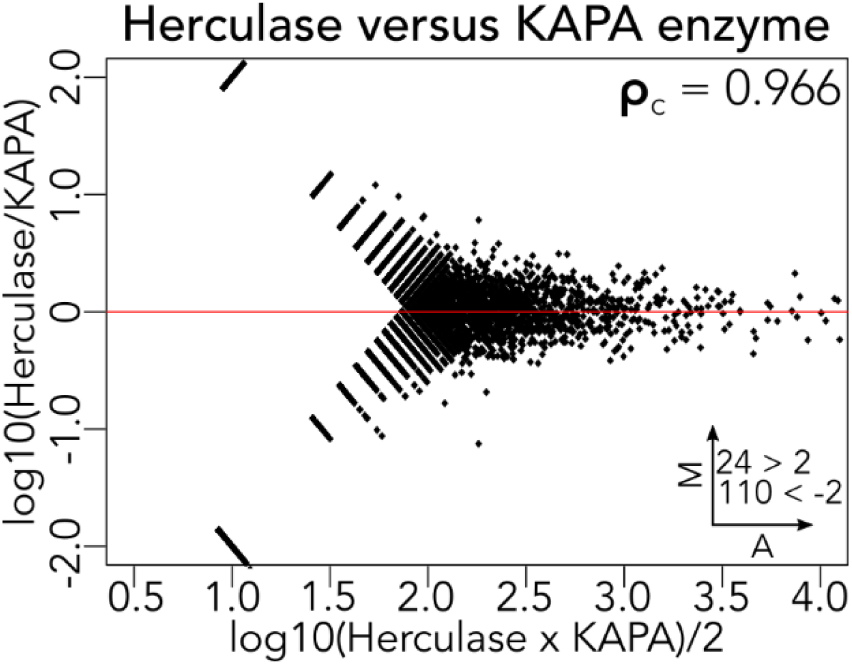
MA plot demonstrates that Herculase II Fusion DNA polymerase performs comparably to KAPA 2G Fast Multiplex Mix in PCR1. CDR3 sequences were included in this analysis if they had a clone count of ≥10 in either data set.

## SUPPLEMENTARY TABLES

**Table S1:**
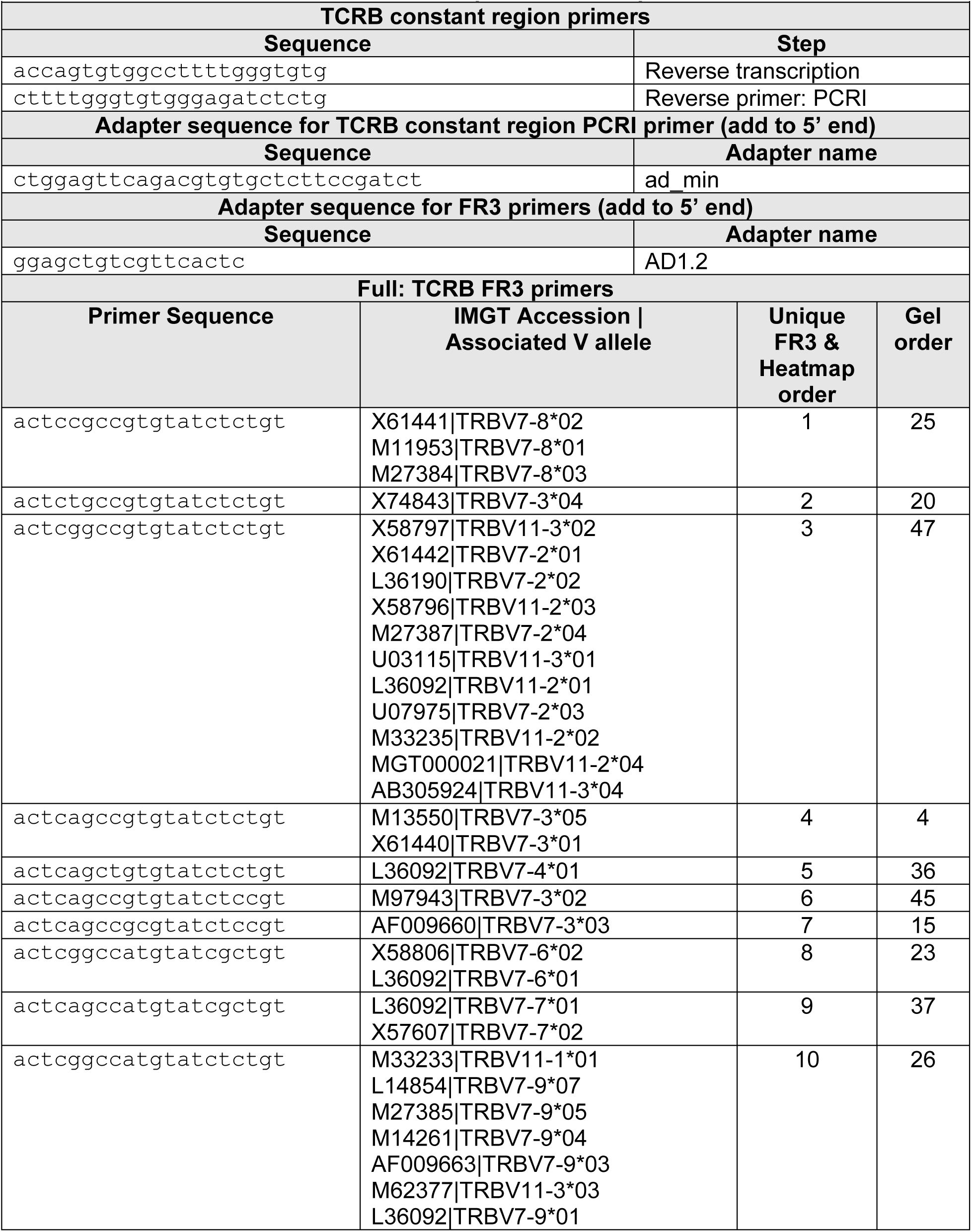
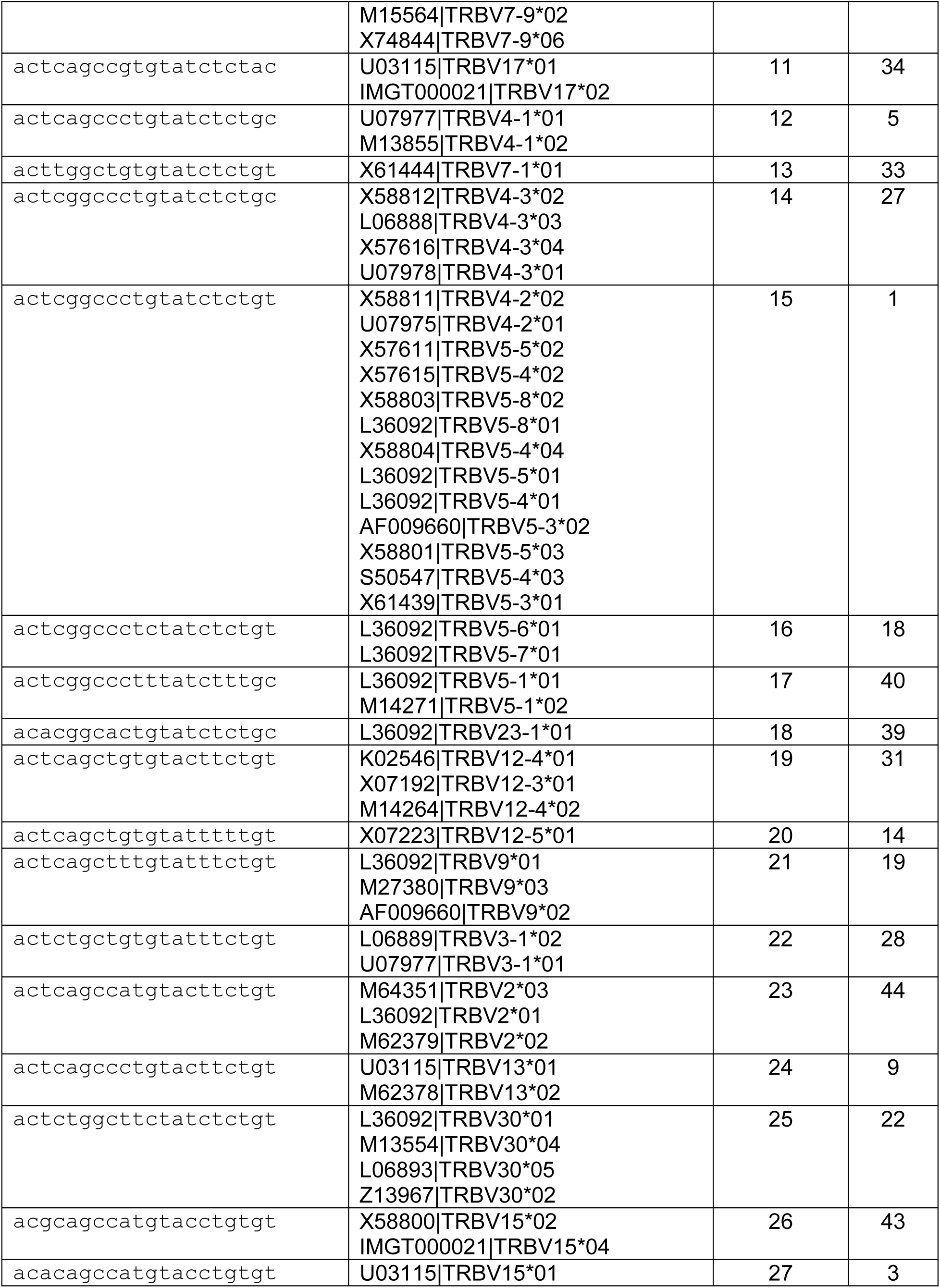
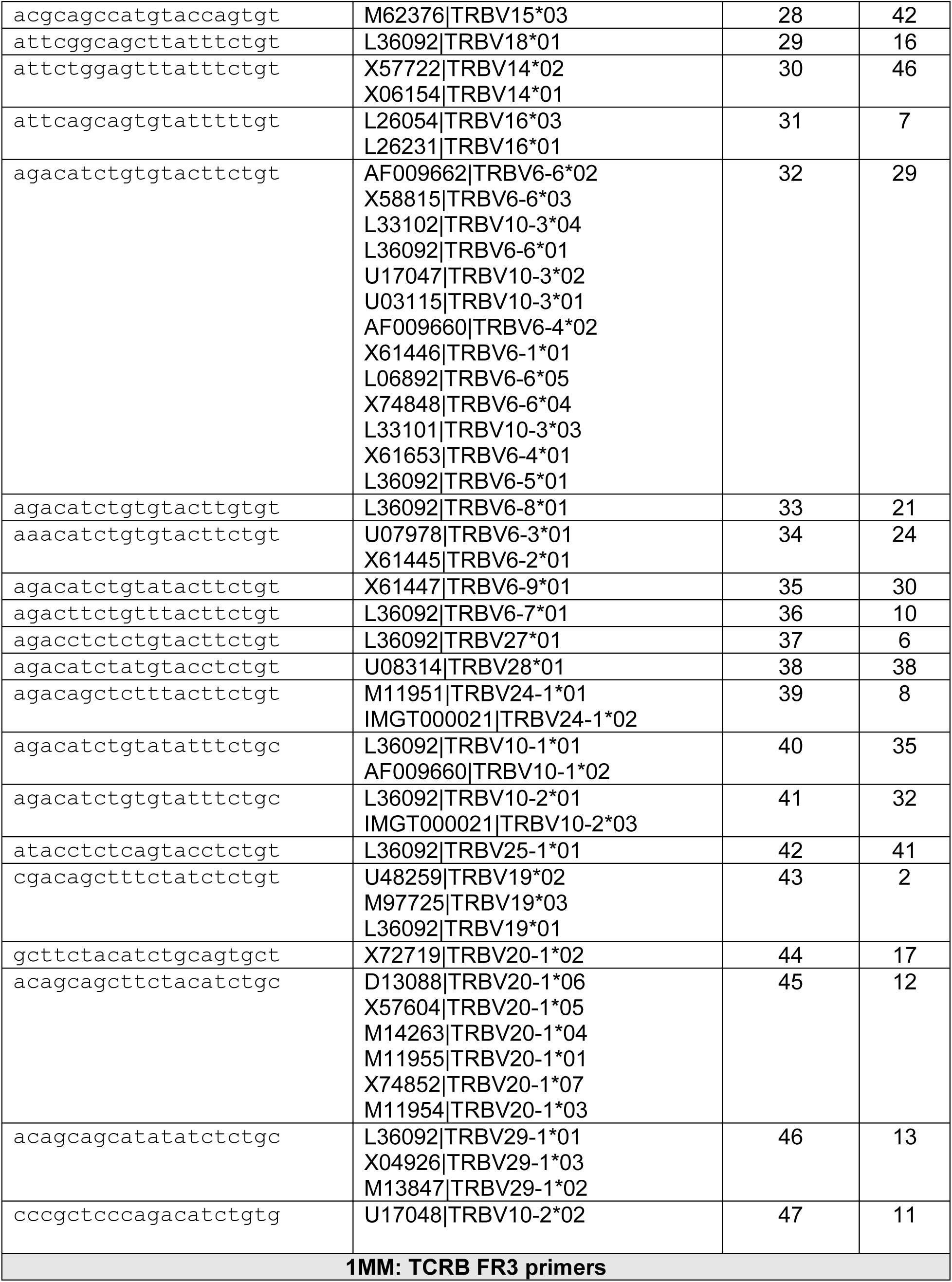
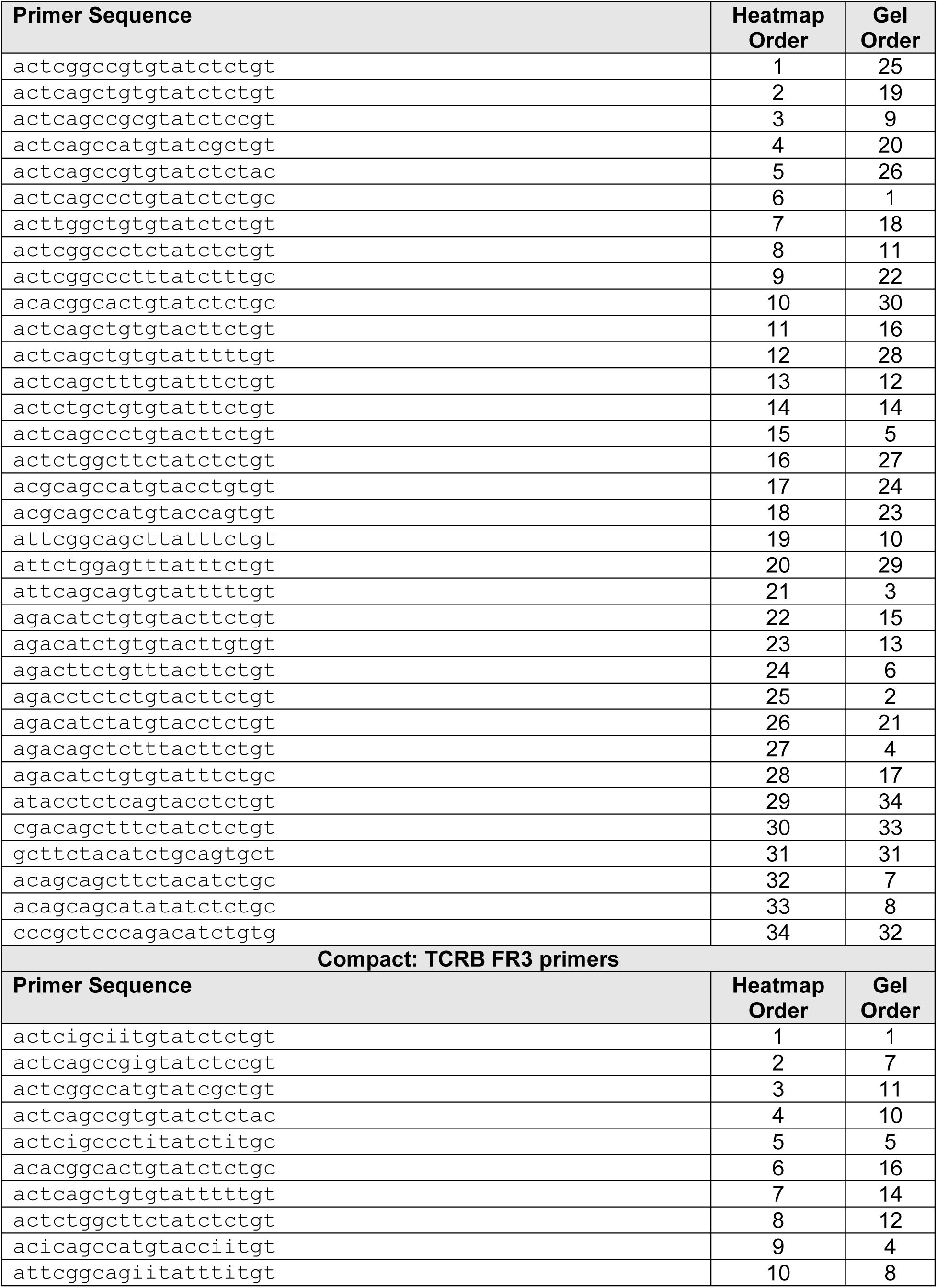
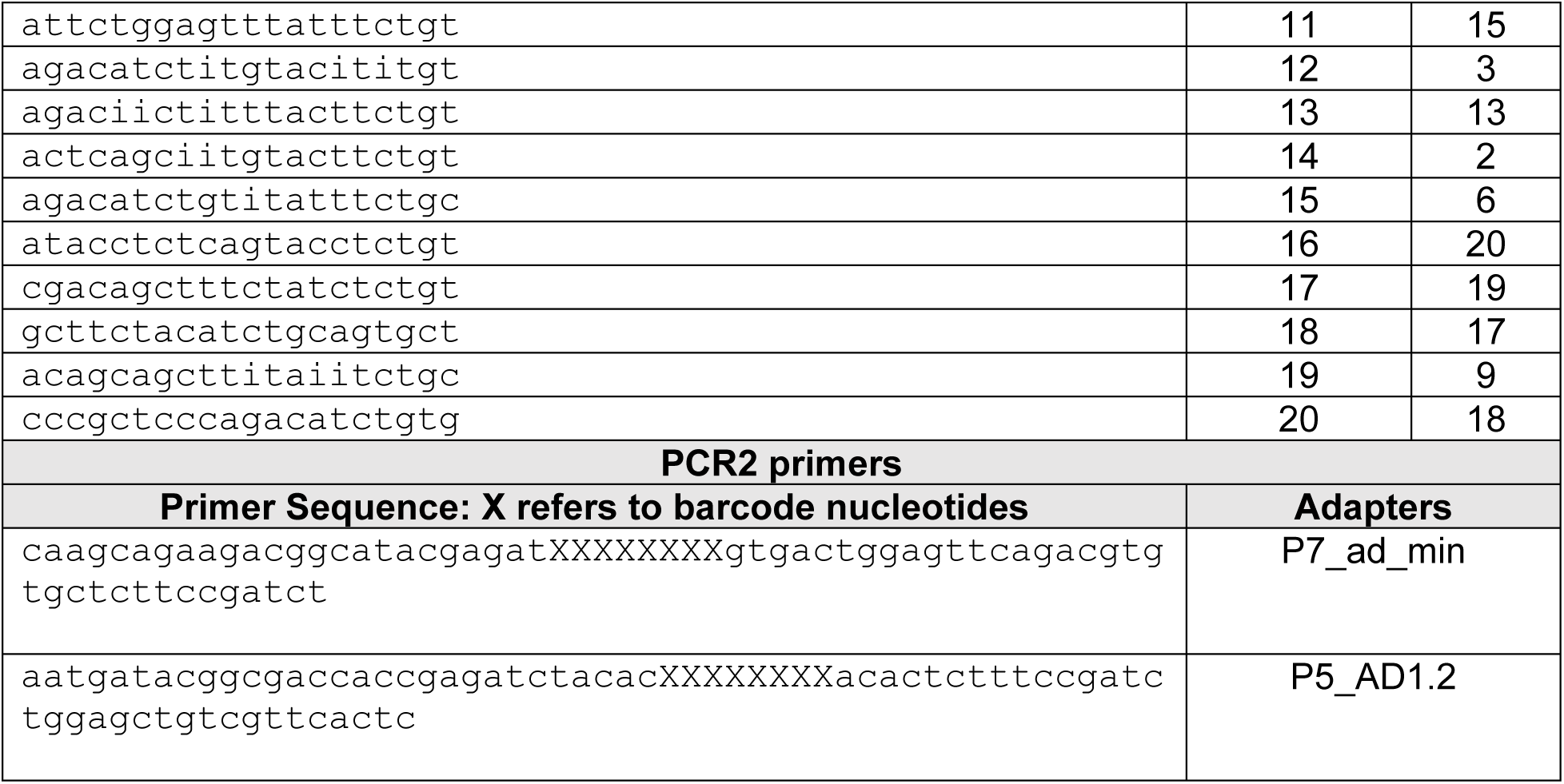
FR3AK-seq human TCRB primers. Human TCRB FR3AK-seq primers and adapter sequences.

**Table S2:**
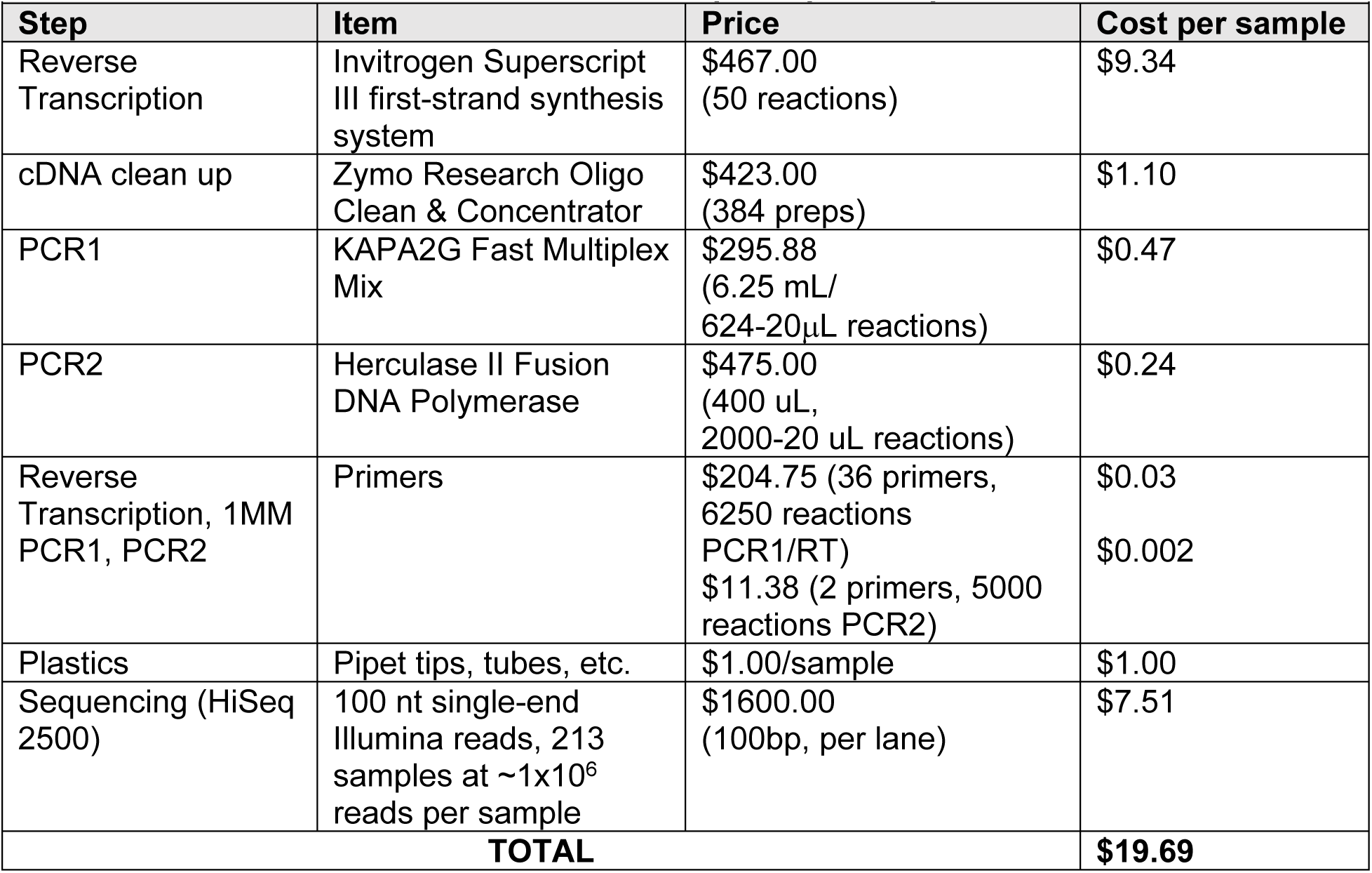
FR3AK-seq cost per sample. Cost per sample for FR3AK-seq. HiSeq cost calculations were based on the Johns Hopkins Genetics Resources Core Facility High Throughput Sequencing Center (internal pricing) at an estimated 1×10^6^ reads per sample. Note that these calculations do not include overhead, personnel costs, or time spent by the researcher.

| **Table S2: FR3AK-seq cost per sample** |  |  |  |
| --- | --- | --- | --- |
| **Step** | **Item** | **Price** | **Cost per sample** |
| Reverse Transcription | Invitrogen Superscript III first-strand synthesis system | \$467.00 (50 reactions) | \$9.34 |
| cDNA clean up | Zymo Research Oligo Clean & Concentrator | \$423.00 (384 preps) | \$1.10 |
| PCR1 | KAPA2G Fast Multiplex Mix | \$295.88 (6.25 mL/ 624-20µL reactions) | \$0.47 |
| PCR2 | Herculase II Fusion DNA Polymerase | \$475.00 (400 uL, 2000-20 uL reactions) | \$0.24 |
| Reverse Transcription, 1MM PCR1, PCR2 | Primers | \$204.75 (36 primers, 6250 reactions PCR1/RT)<br>\$11.38 (2 primers, 5000 reactions PCR2) | \$0.03<br>\$0.002 |
| Plastics | Pipet tips, tubes, etc. | \$1.00/sample | \$1.00 |
| Sequencing (HiSeq 2500) | 100 nt single-end Illumina reads, 213 samples at ~1x10<sup>6</sup> reads per sample | \$1600.00 (100bp, per lane) | \$7.51 |
| **TOTAL** | | | **\$19.69** |

**Table S3:**
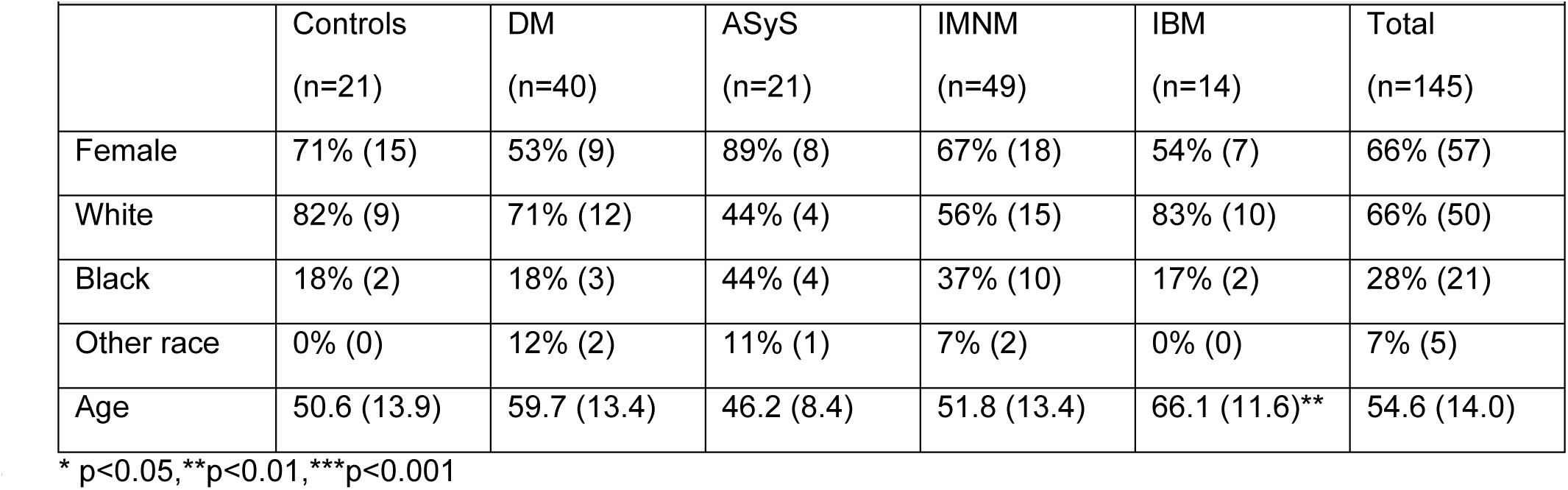
Idiopathic Inflammatory Myopathy and Control Muscle Biopsy Cohort. Description of the IIM muscle biopsy cohort. Chi-squared or Fisheŕs exact tests were used to compare the dichotomous variables of each one of the clinical groups with the rest. Student’s T test was used to compare the age of each one of the clinical groups with the rest.

**Table S4:** GLIPH summary. Summary of IIM and control muscle biopsy FR3AK-seq data used in GLIPH analysis. Bystander clones refer to those appearing at a frequency of less than or equal to 1 cellular equivalent.

| **Table S4: GLIPH summary** |  |
| --- | --- |
| **Input IIM FR3AK-seq data** |  |
| Number of samples with non-bystander T cell clones | 122 (23 excluded) |
| Number of non-bystander T cell clones included in analysis | 11351 |
| Number of bystander T cell clones | 278121 |
| **GLIPH Output** |  |
| Total number of GLIPH clusters | 527 |
| Total number of GLIPH clusters with at least 3 individuals contributing a T cell clone | 143 |
| Number of non-bystander T cell clones clustered by GLIPH | 2125 |

**Table S5:**
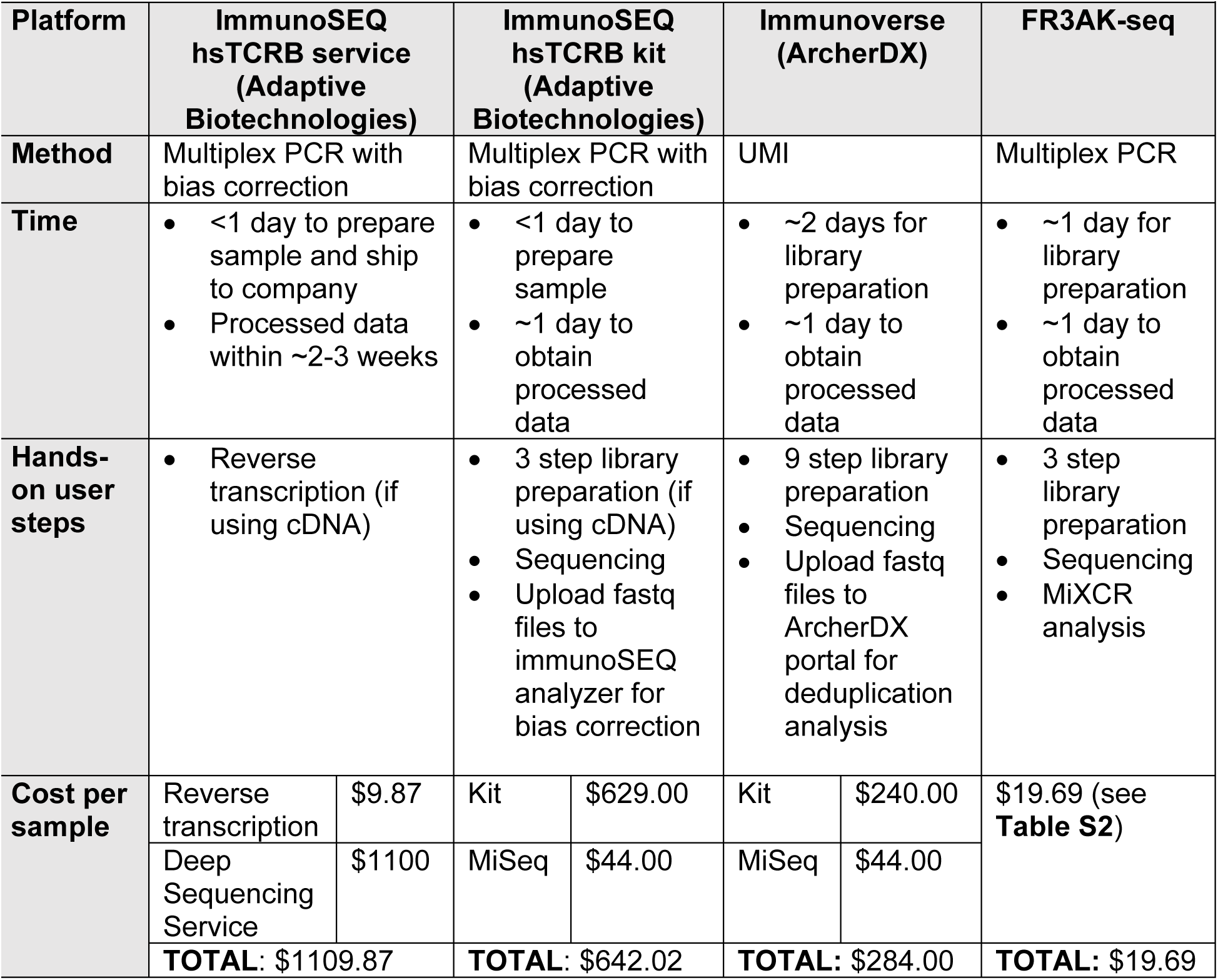
Comparisons between platforms. Comparisons between Adaptive Biotechnologies’ immunoSEQ assay (both service and kit), ArcherDX’s Immunoverse assay, and FR3AK-seq. All prices are approximate. MiSeq cost calculations were based on the Johns Hopkins Transcriptomics and Microarray Core (JHU internal pricing) at an estimated 1×10^6^ reads per sample.

| Table S5: Comparisons between platforms |  |  |  |  |  |  |  |
| --- | --- | --- | --- | --- | --- | --- | --- |
| Platform | ImmunoSEQ<br>hsTCRB service<br>(Adaptive<br>Biotechnologies) |  | ImmunoSEQ<br>hsTCRB kit<br>(Adaptive<br>Biotechnologies) |  | Immunoverse<br>(ArcherDX) |  | FR3AK-seq |
| Method | Multiplex PCR with<br>bias correction |  | Multiplex PCR with<br>bias correction |  | UMI |  | Multiplex PCR |
| Time | <1 day to prepare sample and ship to company<br>Processed data within ~2-3 weeks |  | <1 day to prepare sample<br>~1 day to obtain processed data |  | ~2 days for library preparation<br>~1 day to obtain processed data |  | ~1 day for library preparation<br>~1 day to obtain processed data |
| Hands-on user steps | Reverse transcription (if using cDNA) |  | 3 step library preparation (if using cDNA)<br>Sequencing<br>Upload fastq files to immunoSEQ analyzer for bias correction |  | 9 step library preparation<br>Sequencing<br>Upload fastq files to ArcherDX portal for deduplication analysis |  | 3 step library preparation<br>Sequencing<br>MiXCR analysis |
| Cost per sample | Reverse transcription | \$9.87 | Kit | \$629.00 | Kit | \$240.00 | \$19.69 (see Table S2) |
| | Deep Sequencing Service | \$1100 | MiSeq | \$44.00 | MiSeq | \$44.00 | |
| | **TOTAL: \$1109.87** | | **TOTAL: \$642.02** | | **TOTAL: \$284.00** | | **TOTAL: \$19.69** |

**Table S6:**
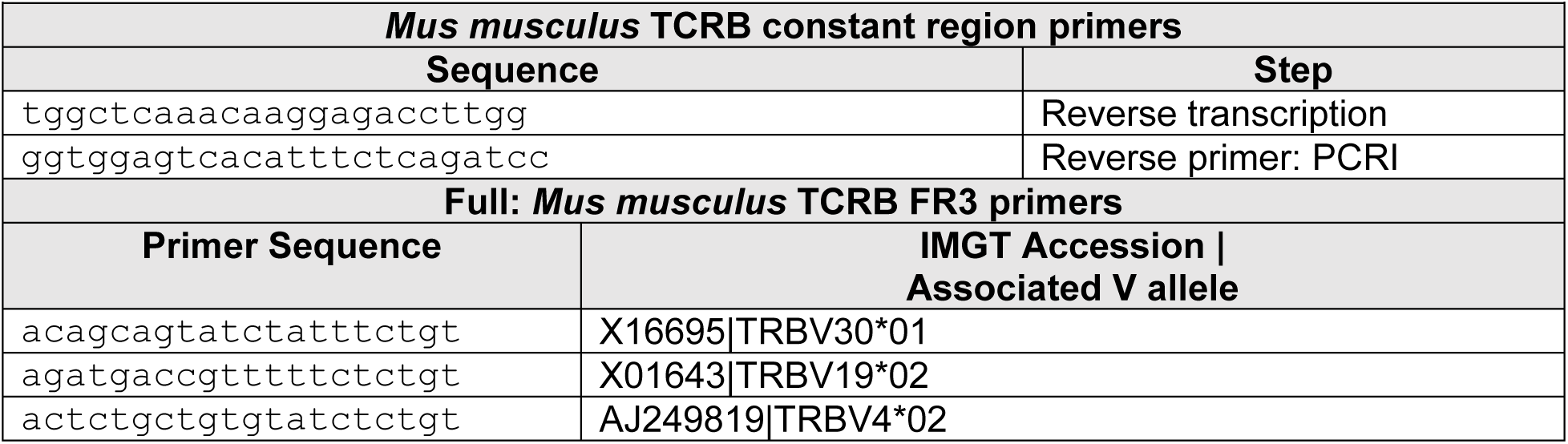
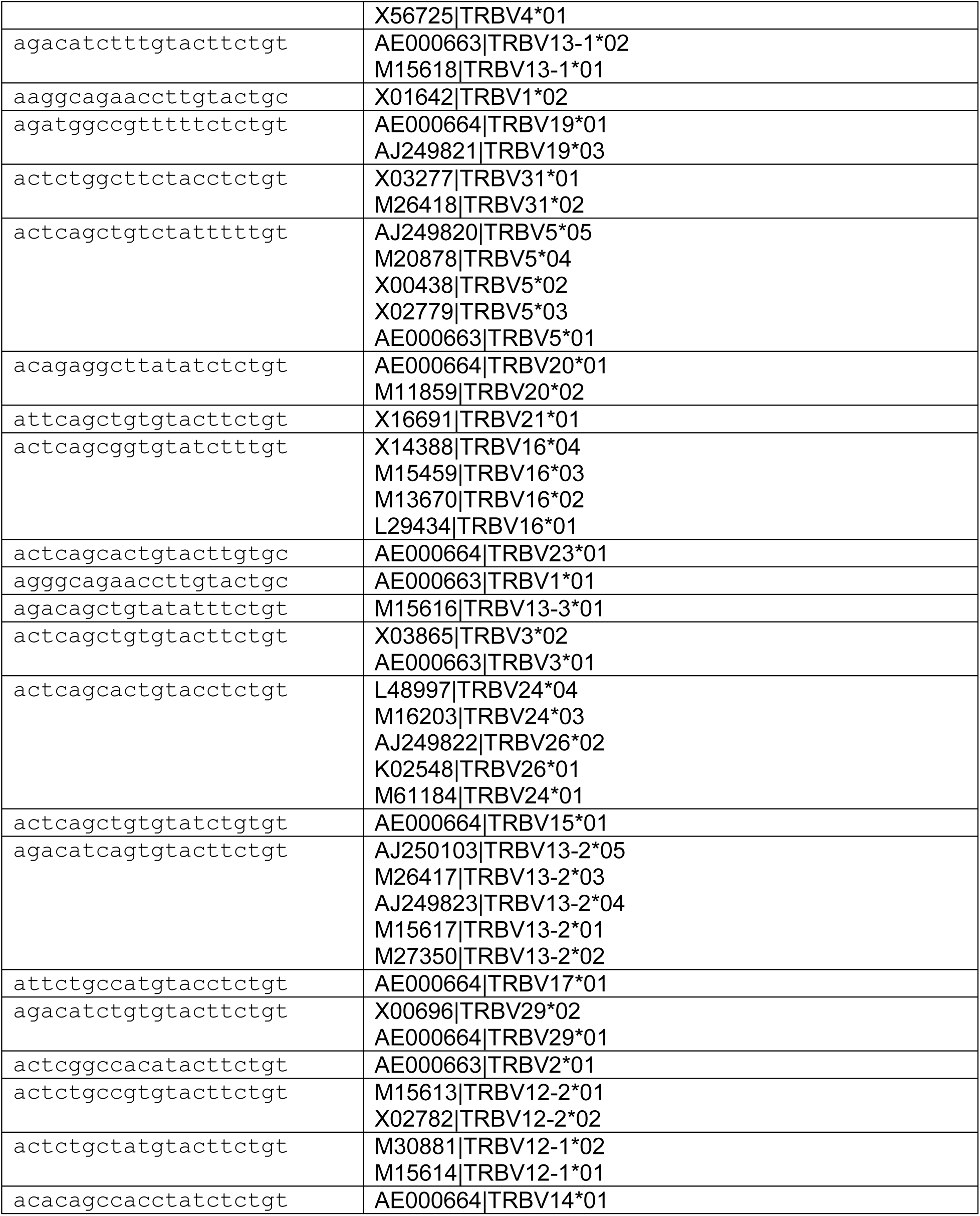
FR3AK-seq Mus musculus TCRB primers. Mus musculus TCRB FR3AK-seq primers. Adapter sequences can be found in Table S1.

**Table S7:**
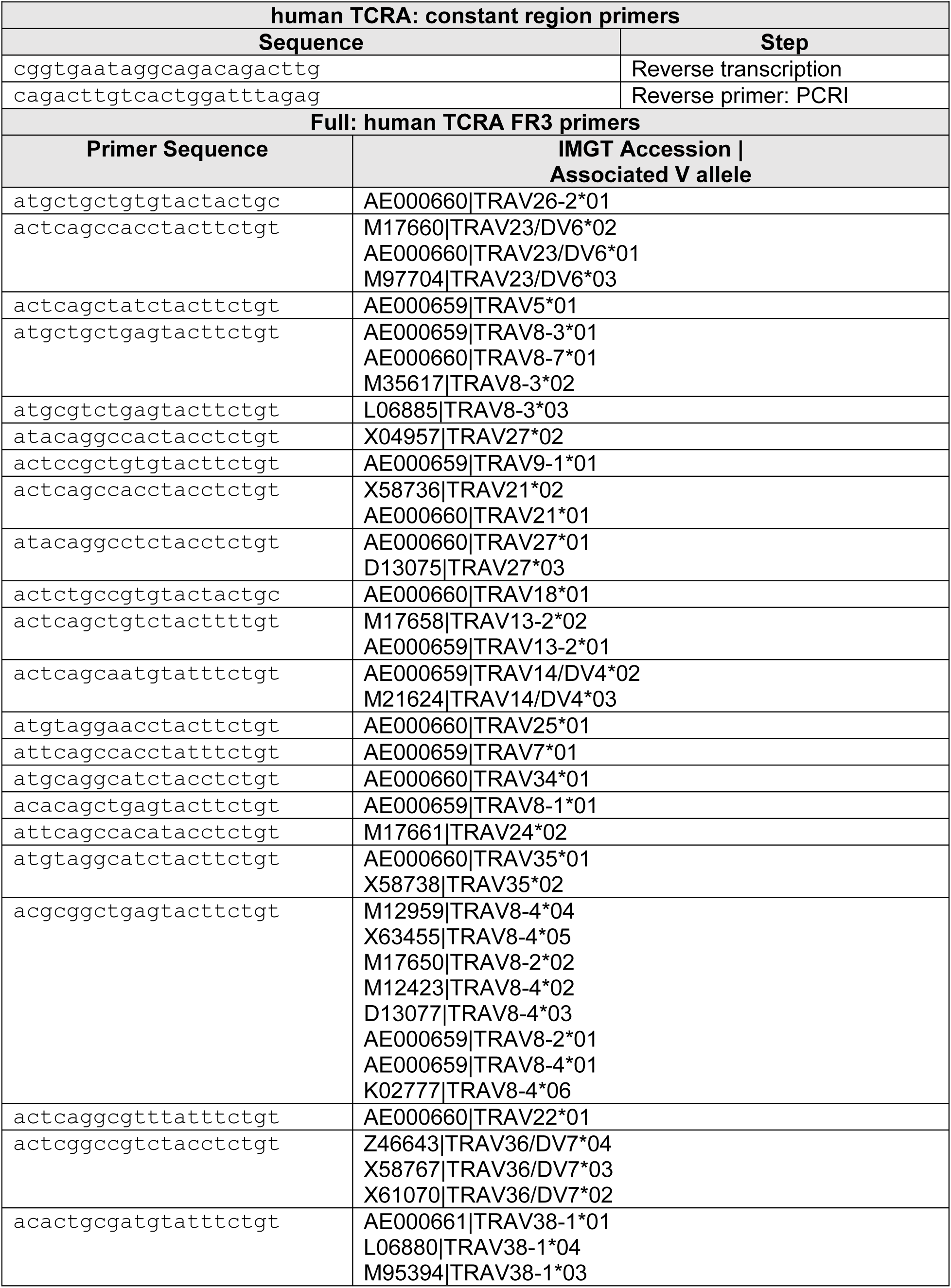
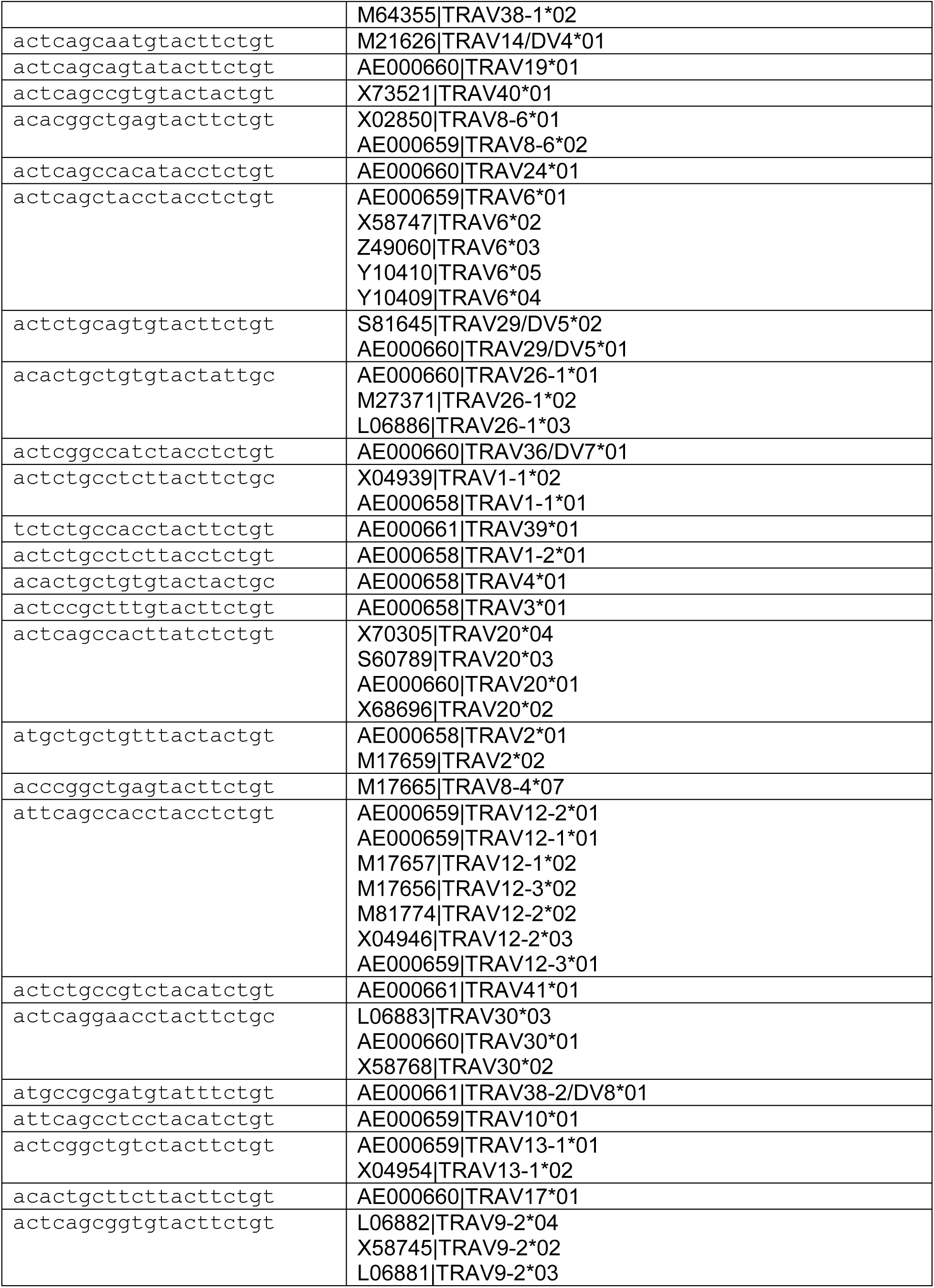
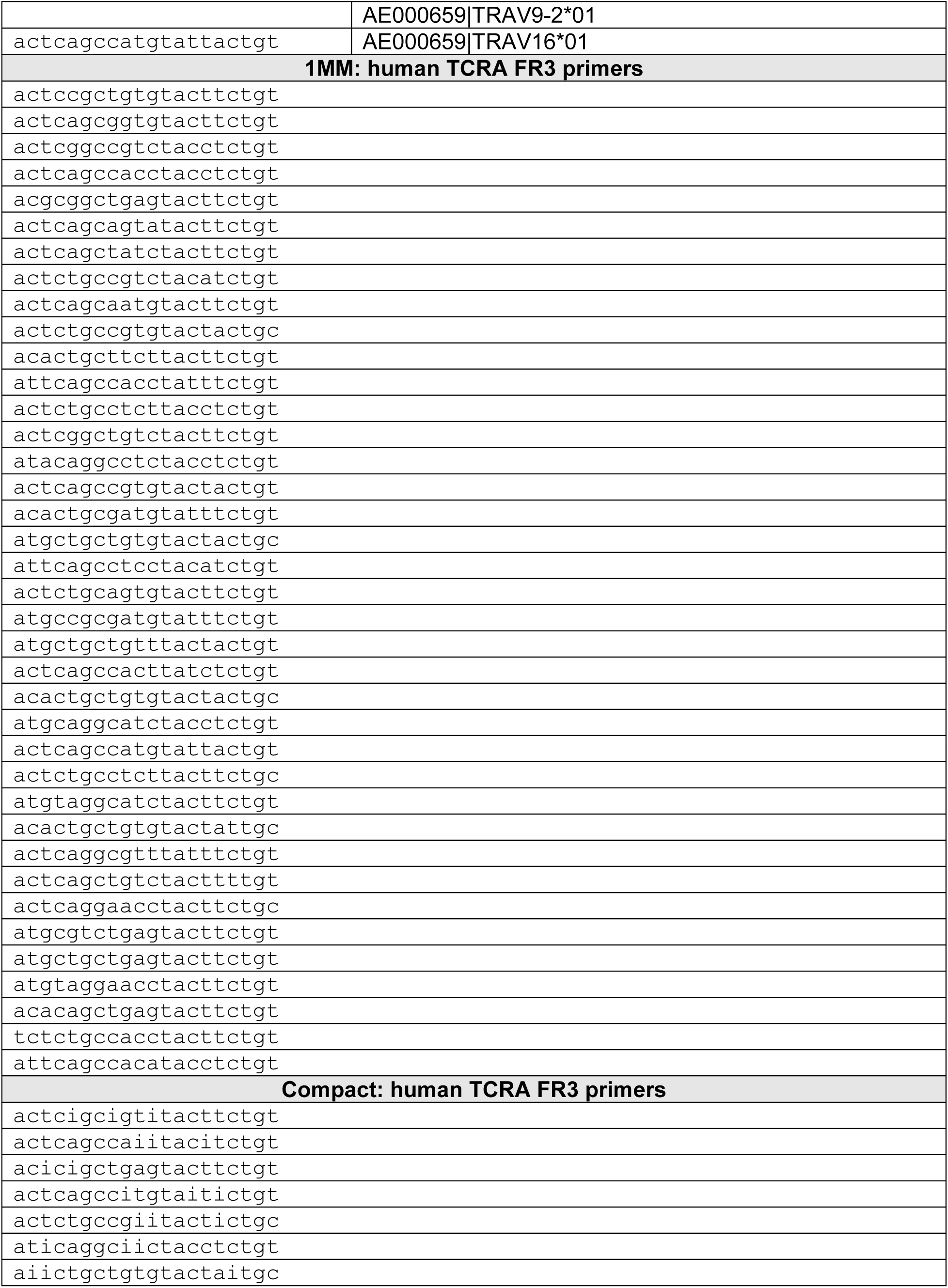
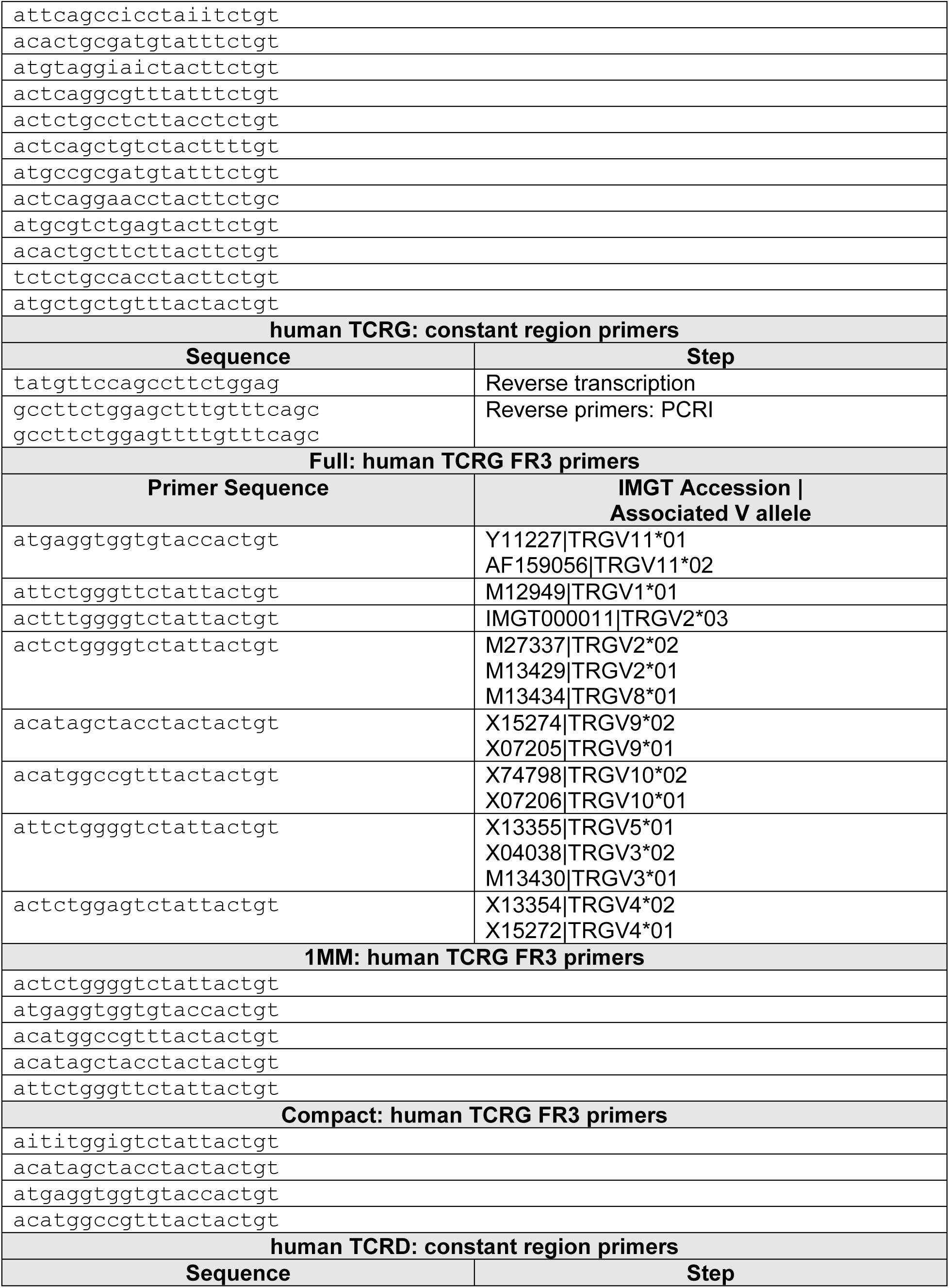
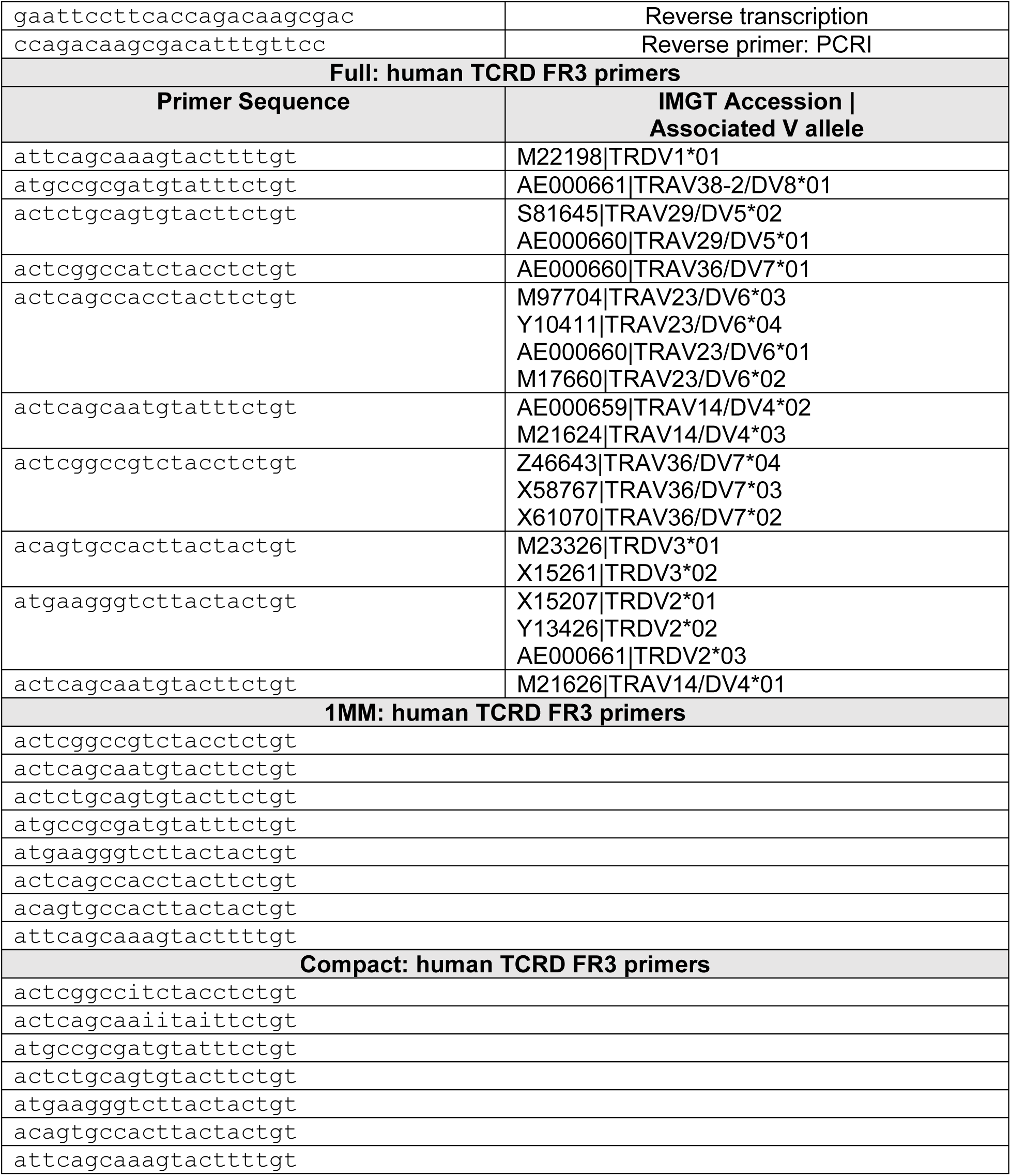
FR3AK-seq human TCRA, TCRG, and TCRD primers. Human TCRA, TCRG, and TCRD FR3AK-seq primers. Adapter sequences can be found in Table S1

**Table S8:**
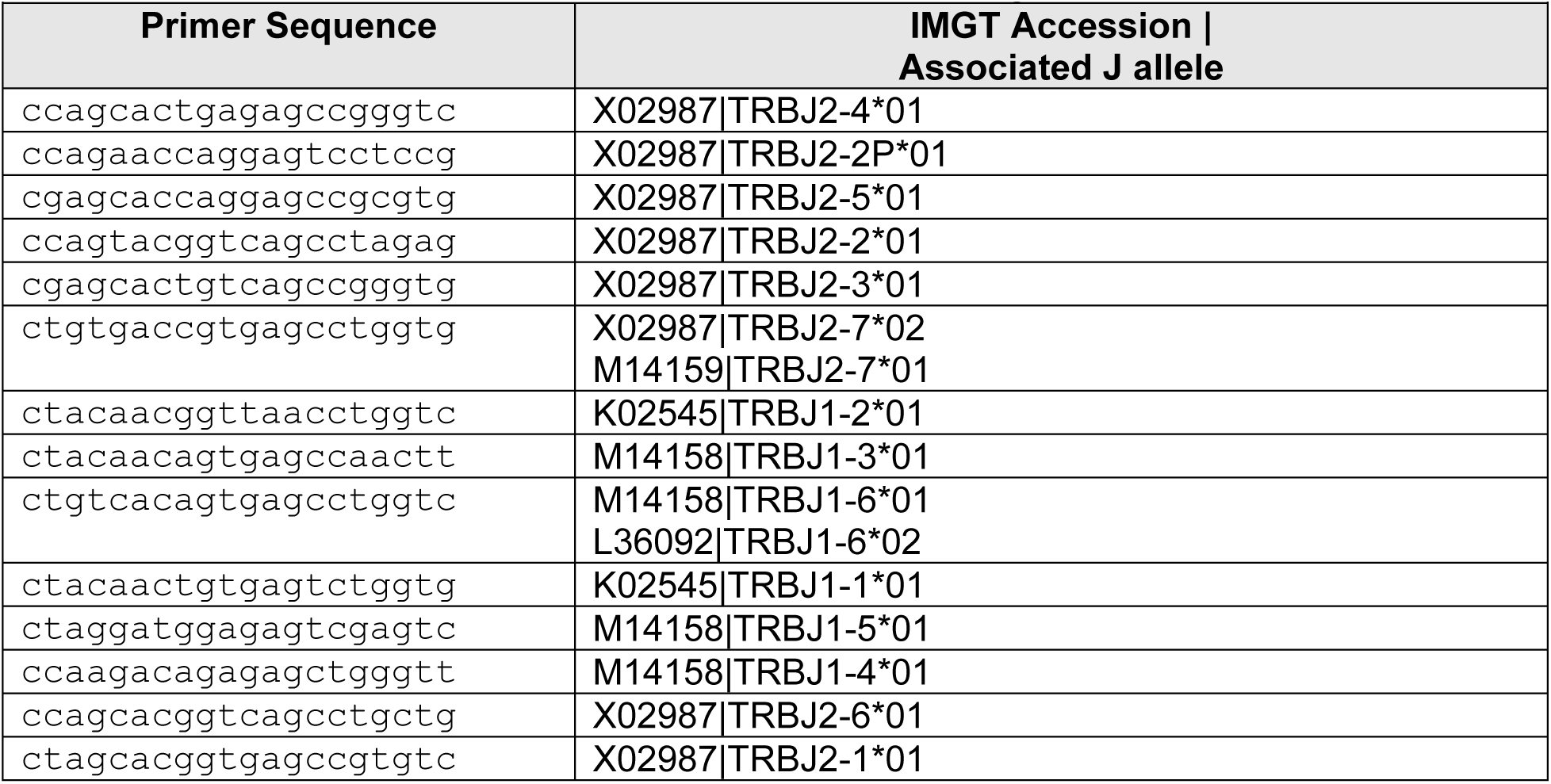
FR3AK-seq human TCRB J region primers. Human TRBJ FR3AK-seq primers. Adapter sequences can be found in Table S1.

**Table S9:** Software parameters. MiXCR and GLIPH parameters used in analysis. Default parameters were used except as noted. Only CDR3b sequences were used with the gliph-group-discovery algorithm. The gliph-group-scoring algorithm was used to match patient and counts with original CDR3b sequences.

| Table S9: Software parameters |  |
| --- | --- |
| MiXCR v2.1.11 |  |
| Parameter | Rationale |
| assemble -OaddReadsCountOnClustering=true | makes error correction algorithms to combine clone abundancies (all “children” clone counts added to head clone) |
| assemble --index | For IIM samples only; links reads in fastq file to CDR3s |
| exportClones --chains TRB | exports TCRB chain sequences only |
| GLIPH 1.0 |  |
| Algorithm step | Parameters |
| --inputs required | Input file(s) column names |
| 1) ./gliph/bin/gliph-group-discovery.pl | default |
| --tcr TCR_TABLE | TCR_TABLE = only CDR3b sequences |
| 2) ./gliph/bin/gliph-group-scoring.pl | default |
| --convergence_file TCR_TABLE-convergence-groups.txt | TCR_TABLE-convergence-groups.txt = from step 1 |
| --clone_annotations FULL_TCR_TABLE | FULL_TCR_TABLE = CDR3b, Patient, Counts |
| --motif_pval_file TCR_TABLE.minp.ove10.txt | TCR_TABLE.minp.ove10.txt Counts = from step 1 |

**Table S10:** Summary of Donor A clones across platforms. Summary of Donor A CDR3 sequences detected across platforms.

| Table S10: Summary of Donor A clones across platforms |  |  |  |
| --- | --- | --- | --- |
| Platform | ImmunoSEQ<br>hsTCRB service<br>(Adaptive<br>Biotechnologies) | Immunoverse<br>(ArcherDX) | FR3AK-seq |
| Total clones detected | 394797 | 67274 | 362288 |
| Number of singletons<br>(clone count = 1) | 345880 | 20570 (pre-deduplication)<br>31668 (post-deduplication) | 198635 |
| Number of clones<br>present at ≥ 10<br>counts | 2702 | 10689 (pre-deduplication)<br>1080 (post-deduplication) | 7189 |

## Notes

#### Summary of Updates

We have added a benchmarking comparison with a UMI-based TCR sequencing assay, expanded our analysis of the quantitative performance of the FR3AK-seq assay, and improved our discussion of the assay and its limitations. We have also removed the part of the paper dealing with mapping of V-alleles.

## References

1. Attaf, M., Huseby, E. & Sewell, A. K. αβ T cell receptors as predictors of health and disease. Cell. Mol. Immunol. 12, 391–399 (2015).

2. Freeman, J. D., Warren, R. L., Webb, J. R., Nelson, B. H. & Holt, R. A. Profiling the T-cell receptor beta-chain repertoire by massively parallel sequencing. Genome Res. 19, 1817– 1824 (2009).

3. Robins, H. S. et al. Comprehensive assessment of T-cell receptor beta-chain diversity in alphabeta T cells. Blood 114, 4099–4107 (2009).

4. Weinstein, J. A., Jiang, N., White, R. A., Fisher, D. S. & Quake, S. R. High-throughput sequencing of the zebrafish antibody repertoire. Science 324, 807–810 (2009).

5. Boyd, S. D. et al. Measurement and clinical monitoring of human lymphocyte clonality by massively parallel VDJ pyrosequencing. Sci. Transl. Med. 1, 12ra23 (2009).

6. Muraro, P. A. et al. T cell repertoire following autologous stem cell transplantation for multiple sclerosis. J. Clin. Invest. 124, 1168–1172 (2014).

7. Gros, A. et al. PD-1 identifies the patient-specific CD8^+^ tumor-reactive repertoire infiltrating human tumors. J. Clin. Invest. 124, 2246–2259 (2014).

8. Beausang, J. F. et al. T cell receptor sequencing of early-stage breast cancer tumors identifies altered clonal structure of the T cell repertoire. Proc. Natl. Acad. Sci. U. S. A. 114, E10409–E10417 (2017).

9. Emerson, R. O. et al. Immunosequencing identifies signatures of cytomegalovirus exposure history and HLA-mediated effects on the T cell repertoire. Nat. Genet. 49, 659–665 (2017).

10. Glanville, J. et al. Identifying specificity groups in the T cell receptor repertoire. Nature 547, 94–98 (2017).

11. Dash, P. et al. Quantifiable predictive features define epitope-specific T cell receptor repertoires. Nature 547, 89–93 (2017).

12. DeWitt, W. S. et al. Human T cell receptor occurrence patterns encode immune history, genetic background, and receptor specificity. eLife 7, (2018).

13. Han, A., Glanville, J., Hansmann, L. & Davis, M. M. Linking T-cell receptor sequence to functional phenotype at the single-cell level. Nat. Biotechnol. 32, 684–692 (2014).

14. Morris, H. et al. Tracking donor-reactive T cells: Evidence for clonal deletion in tolerant kidney transplant patients. Sci. Transl. Med. 7, 272ra10 (2015).

15. Theil, A. et al. T cell receptor repertoires after adoptive transfer of expanded allogeneic regulatory T cells. Clin. Exp. Immunol. 187, 316–324 (2017).

16. Mamedov, I. Z. et al. Quantitative tracking of T cell clones after haematopoietic stem cell transplantation. EMBO Mol. Med. 3, 201–207 (2011).

17. Rosati, E. et al. Overview of methodologies for T-cell receptor repertoire analysis. BMC Biotechnol. 17, 61 (2017).

18. Rudolph, M. G., Stanfield, R. L. & Wilson, I. A. How TCRs bind MHCs, peptides, and coreceptors. Annu. Rev. Immunol. 24, 419–466 (2006).

19. Turner, S. J., Doherty, P. C., McCluskey, J. & Rossjohn, J. Structural determinants of T-cell receptor bias in immunity. Nat. Rev. Immunol. 6, 883–894 (2006).

20. Miles, J. J., Douek, D. C. & Price, D. A. Bias in the αβ T-cell repertoire: implications for disease pathogenesis and vaccination. Immunol. Cell Biol. 89, 375–387 (2011).

21. Kebschull, J. M. & Zador, A. M. Sources of PCR-induced distortions in high-throughput sequencing data sets. Nucleic Acids Res. 43, e143 (2015).

22. Carlson, C. S. et al. Using synthetic templates to design an unbiased multiplex PCR assay. Nat. Commun. 4, 2680 (2013).

23. Shugay, M. et al. Towards error-free profiling of immune repertoires. Nat. Methods 11, 653–655 (2014).

24. Peng, Q., Vijaya Satya, R., Lewis, M., Randad, P. & Wang, Y. Reducing amplification artifacts in high multiplex amplicon sequencing by using molecular barcodes. BMC Genomics 16, 589 (2015).

25. Vander Heiden, J. A. et al. pRESTO: a toolkit for processing high-throughput sequencing raw reads of lymphocyte receptor repertoires. Bioinformatics 30, 1930–1932 (2014).

26. Shugay, M. et al. VDJtools: Unifying Post-analysis of T Cell Receptor Repertoires. PLoS Comput. Biol. 11, (2015).

27. Oakes, T. et al. Quantitative Characterization of the T Cell Receptor Repertoire of Naïve and Memory Subsets Using an Integrated Experimental and Computational Pipeline Which Is Robust, Economical, and Versatile. Front. Immunol. 8, (2017).

28. Ma, K.-Y. et al. Immune Repertoire Sequencing Using Molecular Identifiers Enables Accurate Clonality Discovery and Clone Size Quantification. Front. Immunol. 9, 33 (2018).

29. Egorov, E. S. et al. Quantitative profiling of immune repertoires for minor lymphocyte counts using unique molecular identifiers. J. Immunol. Baltim. Md 1950 194, 6155–6163 (2015).

30. Giudicelli, V., Chaume, D. & Lefranc, M.-P. IMGT/GENE-DB: a comprehensive database for human and mouse immunoglobulin and T cell receptor genes. Nucleic Acids Res. 33, D256–261 (2005).

31. Simsek, M. & Adnan, H. Effect of single mismatches at 3’-end of primers on polymerase chain reaction. J. Sci. Res. Med. Sci. 2, 11–14 (2000).

32. Wu, J.-H., Hong, P.-Y. & Liu, W.-T. Quantitative effects of position and type of single mismatch on single base primer extension. J. Microbiol. Methods 77, 267–275 (2009).

33. Wright, E. S. et al. Exploiting extension bias in polymerase chain reaction to improve primer specificity in ensembles of nearly identical DNA templates. Environ. Microbiol. 16, 1354– 1365 (2014).

34. Ishii, K. & Fukui, M. Optimization of annealing temperature to reduce bias caused by a primer mismatch in multitemplate PCR. Appl. Environ. Microbiol. 67, 3753–3755 (2001).

35. Bolotin, D. A. et al. MiXCR: software for comprehensive adaptive immunity profiling. Nat. Methods 12, 380–381 (2015).

36. Deng, L. et al. Structural basis for the recognition of mutant self by a tumor-specific, MHC class II-restricted T cell receptor. Nat. Immunol. 8, 398–408 (2007).

37. Arahata, K. & Engel, A. G. Monoclonal antibody analysis of mononuclear cells in myopathies. I: Quantitation of subsets according to diagnosis and sites of accumulation and demonstration and counts of muscle fibers invaded by T cells. Ann. Neurol. 16, 193–208 (1984).

38. Fyhr, I. M., Moslemi, A. R., Lindberg, C. & Oldfors, A. T cell receptor beta-chain repertoire in inclusion body myositis. J. Neuroimmunol. 91, 129–134 (1998).

39. Cavazzana, I., Fredi, M., Selmi, C., Tincani, A. & Franceschini, F. The Clinical and Histological Spectrum of Idiopathic Inflammatory Myopathies. Clin. Rev. Allergy Immunol. 52, 88–98 (2017).

40. van Dongen, J. J. M. et al. Design and standardization of PCR primers and protocols for detection of clonal immunoglobulin and T-cell receptor gene recombinations in suspect lymphoproliferations: Report of the BIOMED-2 Concerted Action BMH4-CT98-3936. Leukemia 17, 2257–2317 (2003).

41. Brüggemann, M. et al. Standardized next-generation sequencing of immunoglobulin and T-cell receptor gene recombinations for MRD marker identification in acute lymphoblastic leukaemia; a EuroClonality-NGS validation study. Leukemia 33, 2241–2253 (2019).

42. Greenberg, S. A., Pinkus, J. L., Amato, A. A., Kristensen, T. & Dorfman, D. M. Association of inclusion body myositis with T cell large granular lymphocytic leukaemia. Brain 139, 1348–1360 (2016).

43. Greenberg, S. A. et al. Highly differentiated cytotoxic T cells in inclusion body myositis. Brain J. Neurol. (2019) doi:10.1093/brain/awz207.

44. Bodenhofer, U., Bonatesta, E., Horejš-Kainrath, C. & Hochreiter, S. msa: an R package for multiple sequence alignment. Bioinforma. Oxf. Engl. 31, 3997–3999 (2015).

